# Maps of historical forests in France and their temporal continuity since the first half of the 19^th^ century

**DOI:** 10.64898/2026.07.22.736859

**Authors:** Jean-Luc Dupouey, Laurent Bergès, Nathalie Leroy, Romain Lafite, Frédéric Archaux, Vincent Augé, Raphaël Bec, Maxime Bellifa, Julien Bourguignon, Jérôme Buridant, Béatrice Burlin, Simon Caubet, Albane Chaléat, Sandrine Chauchard, Thomas Cordonnier, Guillaume Decocq, Matthieu Delcamp, Julien Fleury, Sylvain Gaudin, Alain Gervaise, Grégoire Gautier, Julien-Pierre Guilloux, Arnoul Hamel, Wilfried Heintz, Philippe Janssen, Sophie Labonne, Perrine Lair, Thierry Lallemant, Guy Landmann, Laurent Larrieu, Hilaire Martin, Claude Michel, Sylvain Mollier, Christophe Panaïotis, Benoît Renaux, Xavier Rochel, Aline Salvaudon, Marie Thomas, Thierry Touzet, Daniel Vallauri

## Abstract

Integrating environmental history is essential for understanding present-day ecosystem dynamics and guiding modern conservation strategies, particularly under the EU Biodiversity Strategy for 2030. This data paper presents the nationwide digitisation and vectorisation of the French General Ordnance survey map (1818-1866), capturing the country’s forest cover at its historical minimum (a pivotal moment known as the "forest transition").

The original manuscript sheets surveyed at a 1:40,000 scale (976 sheets) were scanned, georeferenced and vectorised. They offer higher thematic accuracy than the older Cassini map and are far more feasible for nationwide vectorisation than the highly detailed Napoleonic cadastre. The area-weighted mean survey date is 1843, and the dataset covers 99.6% of modern mainland France.

To correct for paper deformation, historical surveying errors and coordinate system transformations, a rigorous workflow was established. This shifted from a global 6-parameter affine transformation (root mean square error of 60 m) to a local elastic transformation based on thousands of control points, for 21% of the territory, which reduced the positioning error to 34 m.

We assessed data quality by comparing the General Ordnance Survey maps with the Napoleonic cadastre—the standard reference for 19^th^-century land-use data—across more than 600 municipalities. The correlation between the two sources regarding forest cover percentages was exceptionally high (r>0.9). Because forests formed large, compact blocks of significant strategic interest to military engineers, they were mapped with high precision. Localised inaccuracies were primarily found in remote areas, most notably in the mountains.

The resulting historical vector layer was intersected with contemporary forest data (BD Forêt® v2, 2005-2019). Historical polygons smaller than 0.5 ha were filtered out to comply with modern FAO forest definitions. This spatial overlay generated a new dataset detailing four distinct forest-cover trajectories:

. ancient forests (44.8% of present-day forest): land classified as forest in both the 19^th^ century and the present day (indicating maximum temporal continuity).

. recent forests (55.2% of present-day forest): land that was non-forested in the 19^th^ century but has since undergone reforestation.

. deforested areas (18.9% of 19^th^-century forest): land recorded as forest in the 19^th^ century but subsequently converted to other land uses.

. stable non-forest areas: land that has remained unforested across both periods.

Our results suggest that the forest area of mainland France at its historical minimum should be revised upward to 9.9 million ha. A preliminary analysis further indicates that current spatial variation in forest cover is explained more by land-use dynamics occurring since the forest transition than by the initial extent of forest cover.

These two open-access national datasets (the 19^th^-century forest layer and the land-use transition map) open up new perspectives for a better understanding of present-day forests, including their biodiversity, soil quality, tree growth, and groundwater quality. Furthermore, they provide decision-support tools for conservation planning.

**Key message:** In this data paper, we provide a map of 19^th^-century forests in France based on the digitisation of the military topographic map (1818-1866). By overlaying this historical source with the present-day forest map, we built a second map which allows the identification of ancient forests, recent forests, and deforestation. These maps offer avenues for historical ecology and the design of conservation strategies.

**Links to datasets:** **Map of 19^th^-century forests**

DOI: https://doi.org/10.57745/KSCSQ2

**Map of changes between 19^th^-century forests and present-day forests**

DOI: https://doi.org/10.57745/JZ06QB

## 1 Background

Incorporating history into ecological research is essential to understanding current ecosystem composition, structure and function, as well as defining conservation objectives (Swetnam *et al*. 1999; Jackson and Hobbs 2009; Szabó 2015; Bürgi *et al*. 2017; Bergès and Dupouey 2021; Decoq *et al*. 2022; Navarro *et al*. 2025). Reconstructing long-term temporal trajectories of land use is central to understanding the legacies of former land use on present-day forest ecosystems. This research is however often limited by the availability of land-use data at large spatial and long temporal scales (Santana-Cordero *et al*. 2024). The implementation of early satellite imaging programmes, notably Landsat in 1972 (Wulder *et al*. 2022), has enabled the monitoring of environmental changes on a global scale over the last 50 years. In addition, the declassification of images acquired by American Corona military satellites has made it possible to trace these observations back to the 1960s (Song *et al*. 2014), but not before. Indeed, sources become much scarcer when one attempts to go further back in time and to cover whole countries.

Temporal forest continuity is one of the two components—along with tree and stand age—that define old-growth forests (Cateau *et al*. 2015, Janssen *et al*. 2019). To identify forests with long-term continuity, it is particularly effective to map forest cover at its historical minimum. This point is called the forest transition—the pivot between prolonged deforestation and abrupt, recent reforestation characteristic of industrialised countries (Mather 1992). Forest patches persisting through this bottleneck are likely to have existed long before it. In France, this minimum was reached during the first half of the 19^th^ century. Three major cartographic sources providing national-scale coverage bracket this pivotal period: the Cassini map (1747-1798), which predates the minimum, the Napoleonic cadastre (1808-1850; completed later in Corsica and Savoy) and the General Ordnance survey map (*carte d’État-major*, hereafter the ‘Ordnance map’; 1818-1866), which capture its peak and aftermath.

The Cassini map was drawn at a relatively coarse scale (1:86,400) and provides a simplified representation of land use, making it insufficiently accurate for detailed geo-historical research (Vallauri *et al*. 2012). Furthermore, it does not cover two regions of France: Corsica and the Northern Alps. While the Napoleonic cadastre is highly accurate and drawn at a large scale (generally 1:2,500 or 1:5,000), nationwide vectorisation remains unfeasible at present. This is despite recent attempts to automate the process, such as the work conducted on the 1859 historical cadastral maps of the Province of Trento, Italy (Gobbi *et al*. 2019). Consequently, the Ordnance map, surveyed at 1:40,000, serves as a more accessible alternative of sufficient quality for national studies (see Section 3).

Making historical land-use maps available enables research in various fields, from analysing changes in land use since the Industrial Revolution to identifying the time-lagged effects of historical land use on biodiversity and the functioning of current ecosystems (Lindborg and Eriksson 2004; Helm *et al*. 2006; Krauss *et al*. 2010; Surasinghe and Baldwin 2014; Cousins *et al*. 2015; De Keersmaeker *et al*. 2015; Bergès *et al*. 2016; Bergès *et al*. 2017; Burst *et al*. 2017; Thomas *et al*. 2017; Abadie *et al*. 2021; Lalechère *et al*., 2018; Stein *et al*. 2020; Mollier *et al*. 2022a, b, Mollier *et al*. 2024). In the light of the European Union Biodiversity Strategy for 2030, which aims to mitigate climate change and safeguard biodiversity, historical land-use maps represent a valuable decision-making support for natural area managers when prioritising conservation actions (Grabska-Szwagrzyk *et al*. 2024).

This paper provides two national-scale datasets:

1 The forest layer derived from the Ordnance map for mainland France, surveyed between 1818 and 1866;
2 A map of forest-cover trajectories generated by intersecting the historical Ordnance map with contemporary forest data (BD Forêt® v2, 2005-2019). This spatial overlay allows the identification of four distinct temporal dynamics:

a- ancient forests: lands classified as forest on both the Ordnance map and the present-day map;
b- recent forests: non-forested lands on the Ordnance map, but classified as forest on the present-day map;
c- deforested areas: lands classified as forest on the Ordnance map but subsequently converted to other uses;
d- non-forested areas: land that has remained unforested since the mid-19^th^ century.

## 2 Data and methods

### 2.1 The Ordnance map

The Ordnance map is the oldest high-precision cartographic source to cover nearly all of present-day France (99.6%) and to combine an intermediate scale with a detailed depiction of land use. Produced by the War Office for military purposes, the map accurately represents forests due to their strategic significance. The complete production process is described in detail by Berthaut (1898). The original manuscript sheets, at a scale of 1:40,000, were derived directly from field surveys and depicted land use in colour. They are currently held at the French National Institute of Geographic and Forest Information (IGN) in Paris. These manuscript maps were subsequently reduced to 1:80,000 scale to produce the final engraved, black-and-white map. While the standard survey scale was 1:40,000, specific strategic areas were initially mapped at 1:10,000 or 1:20,000 (Berthaut 1898). The 1:40,000 manuscript maps comprise a total of 976 sheets, each covering an area of 32 x 20 km (excluding truncated border and coastal sheets; Fig. 1).

**Fig. 1.**
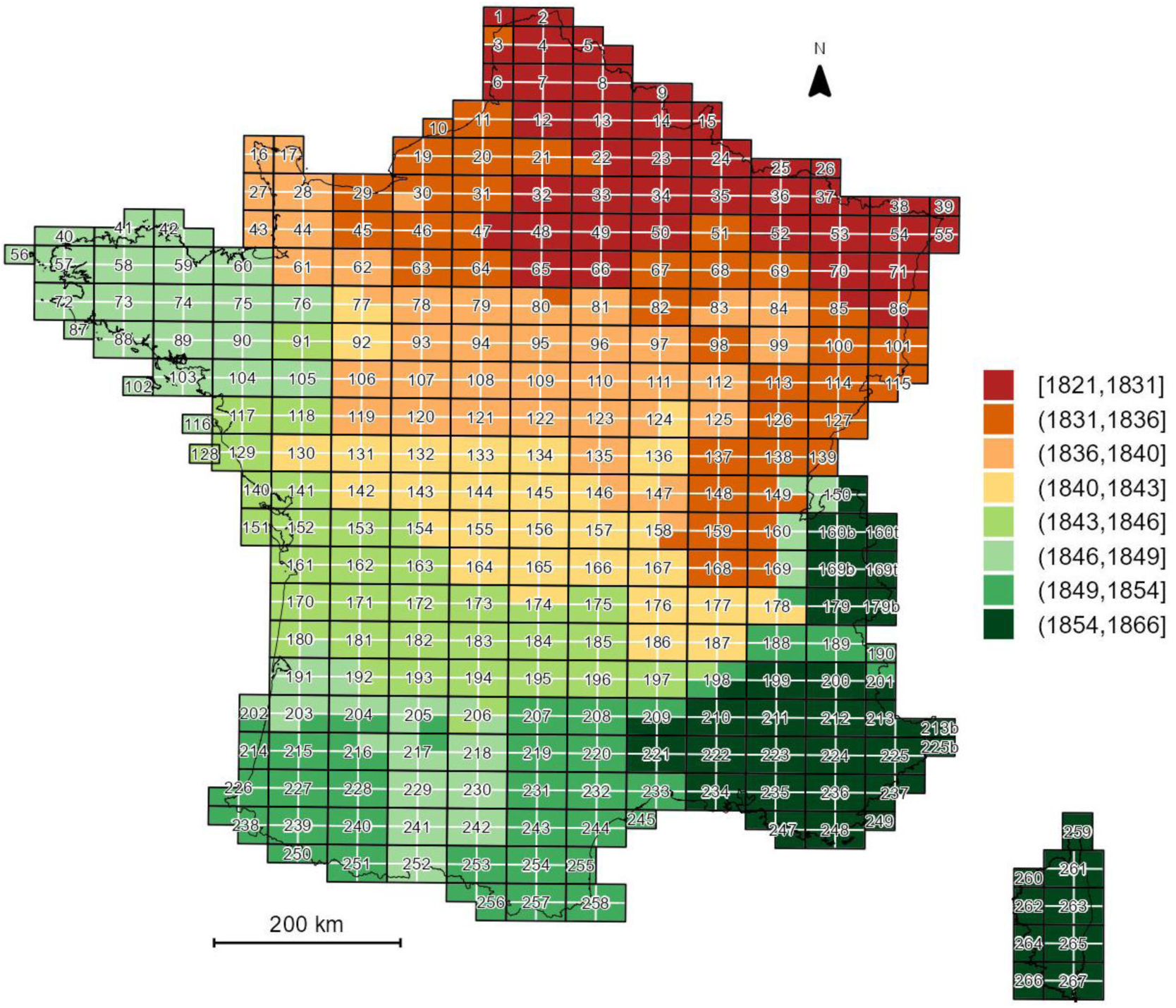
Index map of the 976 original sheets (manuscript maps) of the Ordnance map. The sheets were identified by their number followed by one of the four cardinal directions: NE (north-east), NO (north-west), SE (south-east), and SO (south-west) (e.g., 208NE, 208NO, 208SE, and 208SO). Colours indicate the mean survey date of each sheet.

The surveys were conducted between 1818 and 1866, beginning in the Paris region (the seat of political power) and extending towards the north-eastern border, where major military interests were at stake. They then progressed towards the south-western coast before finally reaching the Alps, Provence, and Corsica (Fig. 1). Savoie, which was incorporated into France in 1860, was the last region to be mapped. The area-weighted mean survey date was 1842.6, while the weighted median was 1842.0, with 75% of the territory surveyed between 1832 and 1852 (Fig. 2). Disruptions in the pace of the surveys coincided with major political regime changes: the 1830 Revolution, the 1848 Revolution, and the advent of the Second Empire in 1852. Regarding land-use representation, the maps relied partly on data from the Napoleonic cadastre. Military geographical engineers used a reduction of the cadastre in the field—most often at a scale of 1:10,000—which they verified and updated as necessary (Rochel *et al*. 2017). It appears that only 6% of the French territory was surveyed *de novo*, independently of the cadastral records (Bacchus and Dupuis 1990).

**Fig. 2.**
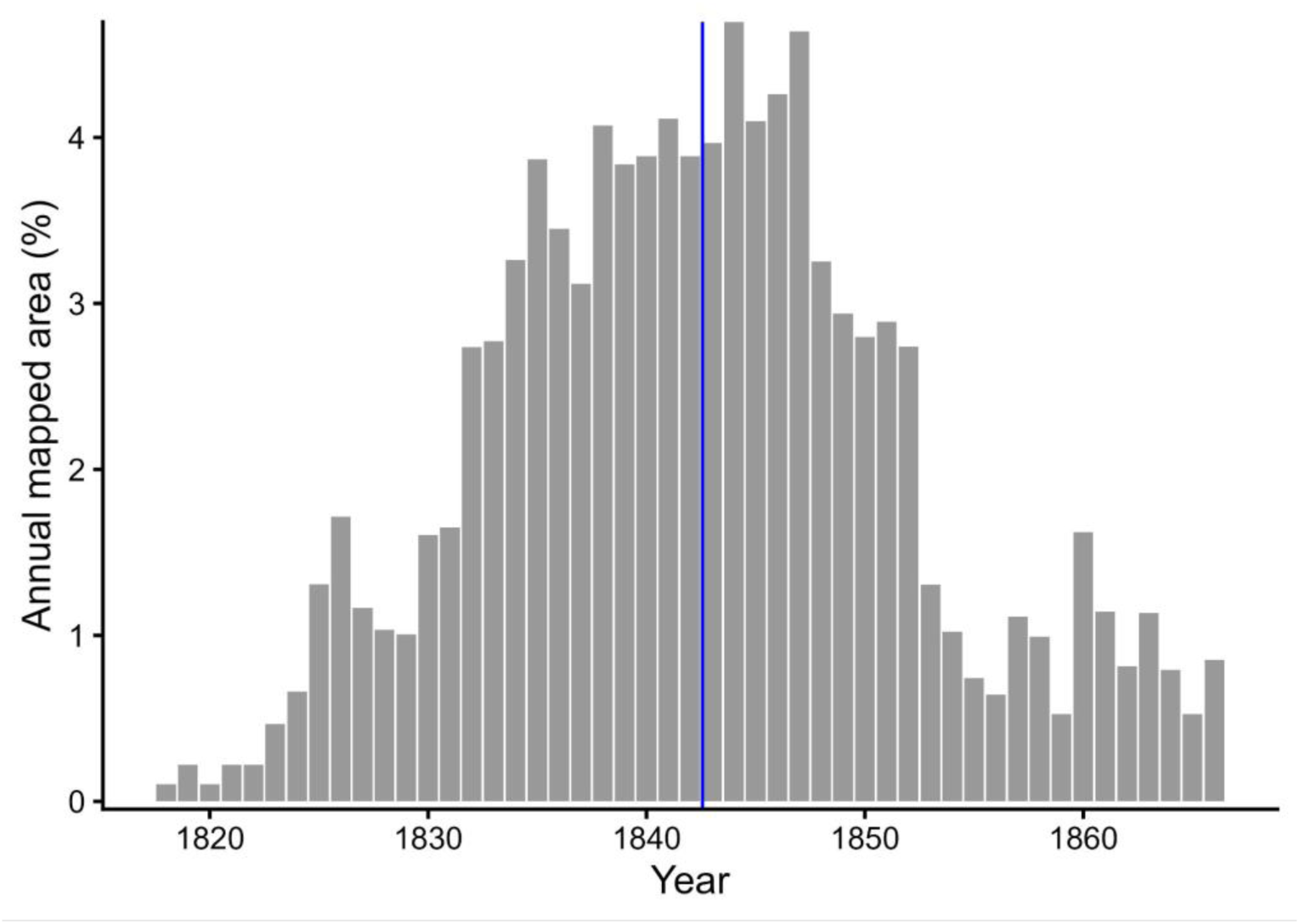
Annual percentage of the total surface area mapped. Since individual map sheets were surveyed over several years, the total area of each sheet (*S*) was distributed equally across the number of years required for its completion (*N*). Thus, for each of those *N* years, the sheet contributed a fractional area of *S*/*N* to the total annual area surveyed across France. The blue vertical line represents the area-weighted mean survey date (1842.6).

Each manuscript map was accompanied by detailed marginal information regarding its creation. One inset map depicted the survey years and the specific areas mapped by each geographical engineer. A second inset (data table) provided the planimetric coordinates and elevations for the third-order triangulation—the densest geodetic level—with an average density of 40 points per sheet (ca. one point per 16 km^2^). The map was based on the Bonne projection, using the Plessis 1817 ellipsoid, with the Paris meridian as the prime meridian and the 45^th^ parallel as the latitude of origin. The Ordnance map does not cover the whole of present-day France, leaving a gap of 193,374 ha across all land use categories (forested or otherwise), which represents 0.4% of the total area of mainland France. This missing area consists of: (a) 70,460 ha annexed under the 1947 Treaty of Paris, located in the former County of Nice (now in the Alpes-Maritimes department; map sheets 213NW, SW & NE, 213bisSW, 225bisNW, and 225NE; Appendix Fig. 3) and in the former Duchy of Savoy (now in the Savoie and Hautes-Alpes departments; sheets 179bisSW, 179SE, 189NE and 201NW); (b) 30,879 ha in Haute-Tarentaise (Savoie department) where only buildings, roads, hydrography, and glaciers were represented, leaving the other land-cover categories unmapped (sheets 169terNW & SW); and (c) the remainder, which stems from discrepancies in the positioning of the coastline within estuaries by historical and contemporary geographers, actual shoreline shifts over the last one and a half centuries, and misalignments of the coastline or land borders resulting from georeferencing errors. These discrepancies also resulted in 52,161 ha being included on the Ordnance map but falling outside present-day France.

Although a commission of experts had defined in detail the different types of land use to be represented on the Ordnance map in 1802, these initial guidelines were only partially followed (Fig. 4). Indeed, numerous variations in the representation of land use, particularly in the use of colours, ultimately appear in the manuscript maps, depending on the choices made by the various cartographers during the 48 years of surveying, though they carry no real thematic significance.

**Fig. 4.**
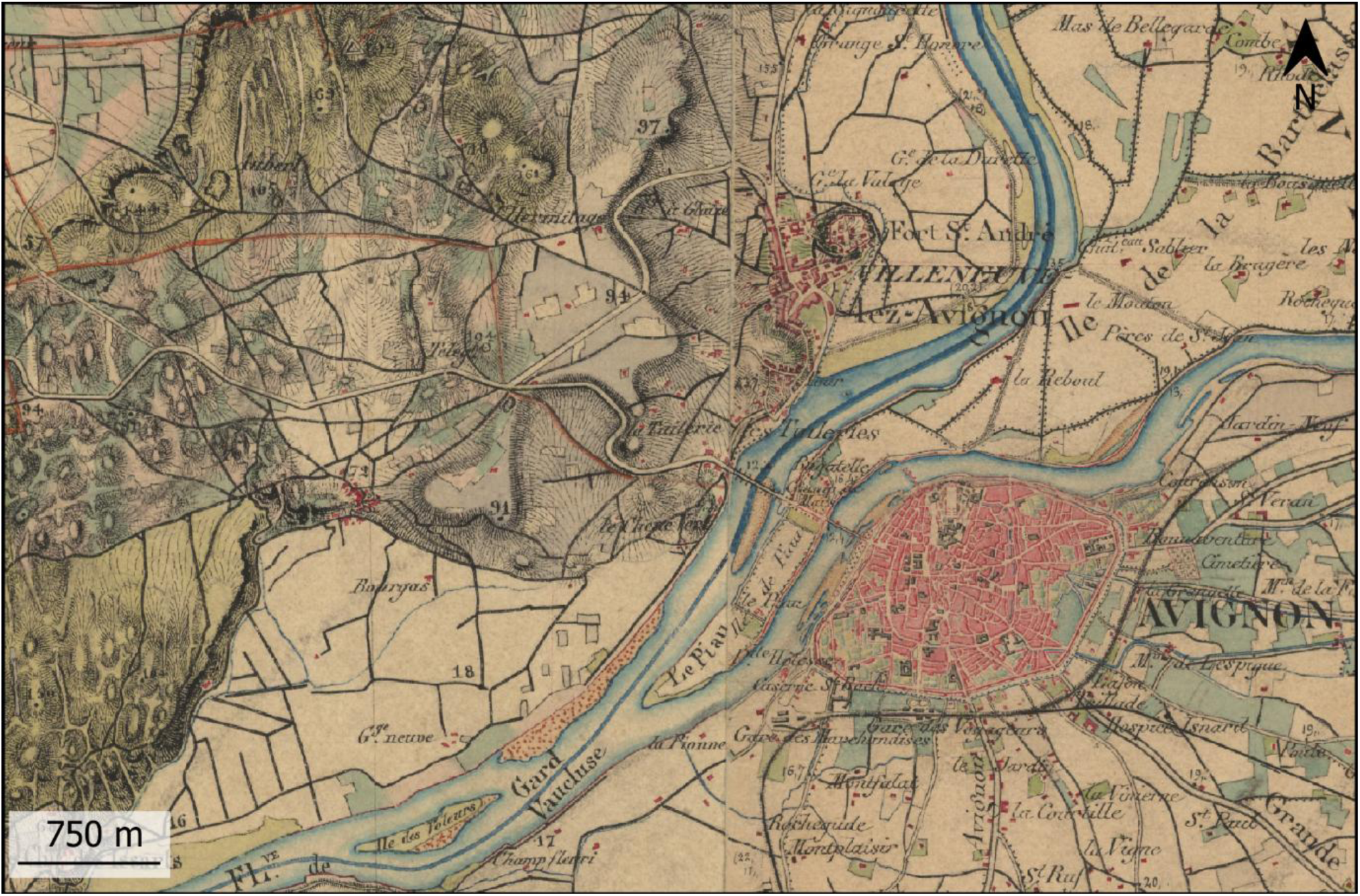
Extract from sheet 222NE (Avignon) illustrating the main land-use categories depicted on the Ordnance manuscript maps. Buildings are shown in red, forests in greenish-yellow, hay meadows in light blue, and vineyards in grey. Pastures are represented by alternating blue and pink washes (north-west), and gravel bars by brownish-orange stippling along the river. Croplands are left unshaded.

Moreover, the commission recommended distinguishing between moorland, heathland, scrubland, wasteland and forests, but we did not find a precise definition of these five related categories. The colours used for moorland, heathland, scrubland and wasteland on the maps were all a mixture of blue and pink, sometimes differing in the shades of blue and pink used.

The land-use classification followed the definitions of the Napoleonic cadastre, where the category was determined by the primary product of the land (*e.g.*, wood production for woodland, grazing for pastures, forage for meadows, crops for croplands, and vines for vineyards). In mountainous areas, the representation of relief using hachures can obscure land-use patterns, making them difficult to identify (Fig. 5A). Representational choices were not limited to semiological issues but also involved the identification and delineation of land use in the field. Fig. 5B illustrates a rare, simplified treatment of land-use boundaries by cartographers, where a sharp, linear break—a seamline artifact—separates the forest on the right from the grassland on the left.

**Fig. 5.**
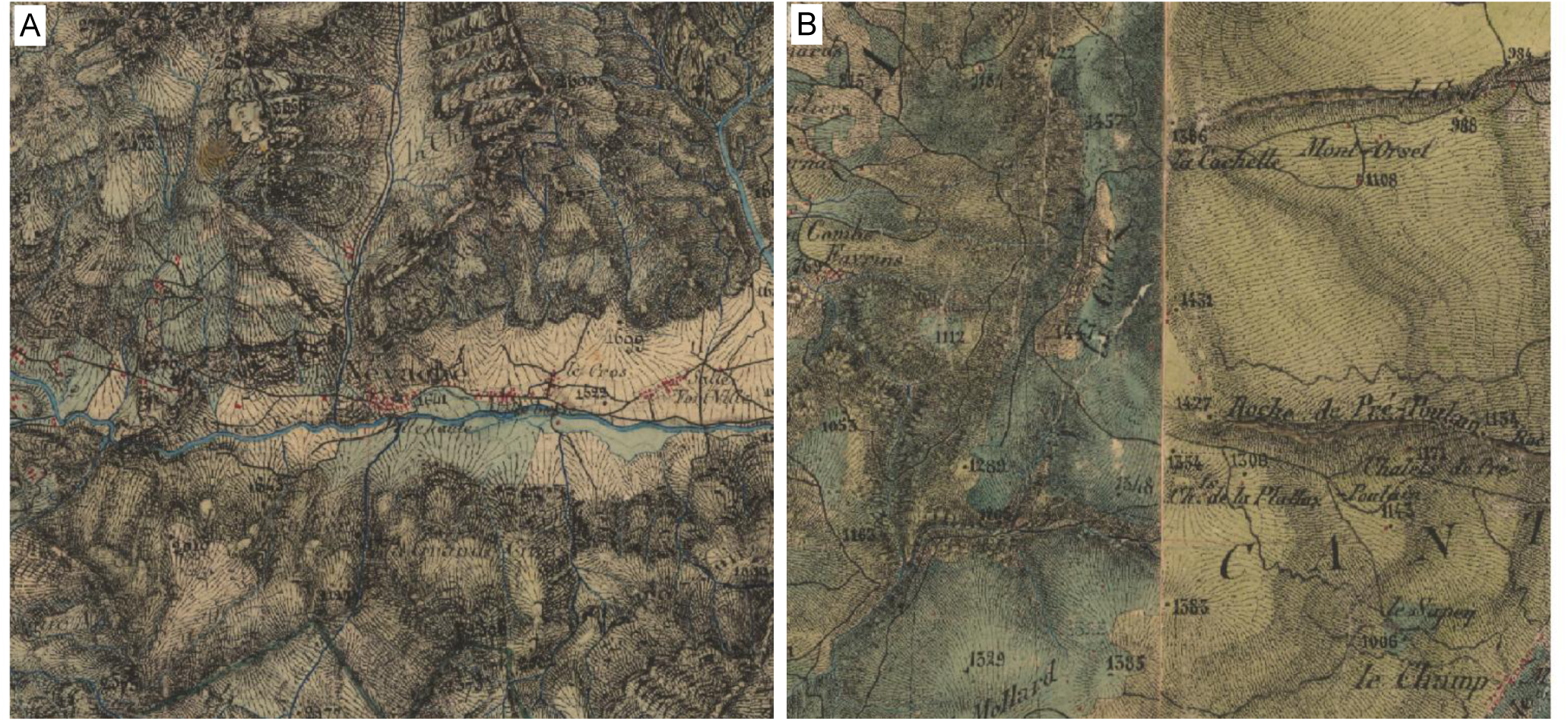
Extracts from the Ordnance manuscript map illustrating two sources of error in the reconstruction of historical land use. Each extract represents a 6 x 6 km area. (A) Névache Valley (Hautes-Alpes department), manuscript map 189NE (Briançon): interpreting land use in mountainous areas is more challenging because it is superimposed on hachures, used to depict the steepness of the terrain. (B) Massif des Bauges (Savoie department), manuscript map 169bisNW (Albertville): in this area, where meadows predominated in the west and forests in the east at the time, according to the Napoleonic cadastre, the boundary between these two zones suffers from an edge-matching issue. Mapped by two different geographers in 1862 and 1864, respectively, the limit is too linear.

Certain symbols were geographically restricted to regions where the corresponding features were common, and were never used elsewhere, despite the probable presence of these features in other areas. For instance, wooded pastures were only mapped as a distinct category in the Jura Mountains.

Eighty-six distinct land-use symbols were identified during the vectorisation of the manuscript maps (Favre *et al*. 2017). The interpretation of these symbols was based on cross-referencing with the Napoleonic cadastre, where it was digitised and available through the Departmental Archives (*e*.*g*., for the Isère department: https://archives.isere.fr/). These symbols were consolidated into a list of 37 classes (Touzet and Lallemant 2016, Lallemant *et al*. 2017). To produce the historical forest map, we selected: the forests (class EM9.1 in Touzet and Lallemant 2016), swamp forests (classes EM9.1 and EM11.1 superimposed), wooded dunes (EM9.1 and EM12.2), wooded rocks and scree (EM9.1 and EM14.1) and castle parks (EM2.2) from these 37 classes.

No minimum size was imposed on military geographers for the polygons they drew. The smallest forest polygons we observed were 150 m² in area. A few polygons in the provided layer, specifically those along the French border, may be even smaller because these were originally larger polygons extending beyond modern-day France which were subsequently clipped by the national border. Some small woodlots may be missing. For instance, a 0.9 ha wood was found to be absent from the map, despite microhistorical research confirming its existence at the time of the survey (Lathuillière & Gironde-Ducher 2021).

### 2.2 Vectorisation

Vectorisation was carried out on scans of the Ordnance manuscript maps at a scale of 1:40,000. These were scanned either at a resolution of 254 dpi (equating to 1 pixel per 4.00 m on the ground) or at 600 dpi (1 pixel per 1.69 m on the ground). Vectorisation was performed manually, with a strict requirement not to deviate from the land-use polygon outlines by more than 2 or 3 pixels. Despite recent progress (Herrault *et al*. 2015; Auffret *et al*. 2017; Gobbi *et al*. 2019; Martinez *et al*. 2023; Mäyrä *et al*. 2023; Ågren and Lin 2024; Andrieu and Serres 2024; De Keersmaeker *et al*. 2026), fully automated approaches to land-use recognition on historical maps remain challenging and have yet to yield satisfactory results. Final error rates remain high—whether in raw AI-generated outputs or after filtering out small polygons—and necessitate extensive manual verification. Only one published study (Herrault 2015) has provided quantitative performance metrics for automated recognition of forests on French Ordnance maps. For three map sheets, the F1-score —a measure of prediction reliability— ranged between 69.5% and 90.6%. An error rate exceeding 30% may be unacceptable to many users of historical maps, especially when working at a local scale.

In 64.5% of the territory, only forests were vectorised, whereas in the remaining 35.5%, all land-use categories (including forests) were identified and vectorised (Table 2). In the dataset provided, only the forest layer is included.

**Table 1.** Georeferencing workflow applied to the Ordnance manuscript maps to produce the vectorised historical forest map of France. RMSE: Root Mean Square Error.

| Level | Coord. system | Transformation method applied to each map sheet | Nature of corrections | RMSE (m) |
| --- | --- | --- | --- | --- |
| 1 | None (image pixel) | None | None | - |
| 2+3 | RGF93 v1 / Lambert-93<br>EPSG 2154 | 2: Polynomial transformation based on 38 points per sheet (5.9 points/100 km<sup>2</sup>) to the ATIG-Bonne (19<sup>th</sup> century) projection<br>3: Three-parameter translation between geocentric cartesian coordinates of the 19<sup>th</sup> century and the current system | Distortions due to scanning, ageing of the paper medium and gluing in the original map | 85<br>(47 to 143) |
| 4 |  | Affine transformation of polygons using a grid of 11,835 control points distributed in the whole of France (2.1 points/100 km<sup>2</sup>) | All types of errors, including errors made during topographic surveying, engraving, and map printing | 60<br>(41 to 95) |
| 5 |  | Elastic transformation of polygons based on 500 to 3,000 control points per sheet (80 to 470 points/100 km<sup>2</sup>) |  | 34<br>(23 to 47) |

**Table 2.** Percentage of total land area (54 773 653 ha) vectorised from the Ordnance map, categorised by georeferencing accuracy (level 4 vs. level 5) and land-use digitisation scope (forests only *vs*. all land-use classes).

|  |  | Georeferencing accuracy |  |  |
| --- | --- | --- | --- | --- |
|  |  | Level 4 | Level 5 | Total |
| Types of land use vectorised | Forests only | 63.9 | 4.1 | 68 |
|  | All land uses (including forests) | 15.2 | 16.8 | 32 |
| Total |  | 79.1 | 20.9 | 100 |

### 2.3 Georeferencing

The georeferencing of the scanned images aimed, firstly, to transform the original coordinates into a modern coordinate system and, secondly, to compensate for positioning errors in the original maps, paper deformation over time and distortions from the scanning process.

Initially, the Ordnance manuscript maps were scanned images without an associated geographical coordinate system (Table 1, level 1). These non-georeferenced sheets are available for download via the IGN website (https://remonterletemps.ign.fr). Coordinates from the 19^th^-century projection system (ATIG-Bonne), indicated in the marginalia of the maps, were assigned to the pixels using 38 control points per sheet and a polynomial transformation (level 2). These coordinates were then transformed from the historical ATIG-Bonne system into the current RGF93 v1 / Lambert-93 coordinate system (level 3). These level-3 georeferenced sheets, organised into 13 regions, are available as a standardised product (SCAN État-major® 40K) through the French government’s geospatial portal (https://cartes.gouv.fr/rechercher-une-donnee/dataset/IGNF_SCAN-EM-40K).

The level-3 sheets were then georeferenced using a non-elastic, 6-parameter affine transformation based on a grid of 11,835 control points distributed as evenly as possible across France (consisting of church steeples, averaging 2.1 points per 100 km²), to produce level-4 maps. For a portion of the territory (20.4%, Table 2 and Appendix Fig. 6), depending on the project sponsor’s requirements and the time available, an elastic georeferencing was performed. This process used a much denser set of control points (500 to 3,000 points per sheet, or 80 to 470 points per 100 km²) to generate level-5 maps, starting either from level-1 or from level-3 sources. This local georeferencing combined a Delaunay triangulation of control points and the 6-parameter affine transformation in each triangle. It relied on a wide variety of landmarks, including buildings (castles, manor houses, mills, isolated farms), religious structures (chapels, cemeteries, crosses, calvaries), and network nodes (hilltops, crossroads, sharp road bends, and hydrographic confluences).

Based on a sample of 28 sheets distributed across France and using a total of nearly 36,000 control points, the root mean square error (RMSE) of positioning was calculated for each sheet. At levels 3 and 4, the RMSE was derived directly from the transformation residuals. Since elastic transformation at level 5 does not produce residuals, the RMSE was determined using a leave-one-out cross-validation procedure repeated for each control point: each point was iteratively removed, the elastic georeferencing recomputed without it, and the error measured at the omitted point. These errors were then aggregated to calculate the sheet-level RMSE. Overall, the mean RMSE values decreased from 85 m at level 3 to 60 m at level 4, and to 34 m at level 5 (Table 1). The maps provided here combine geodata that were georeferenced to level 4 or 5 (Table 2).

### 2.4 Assessing forest dynamics: spatial intersection of the Ordnance map and the present-day forest map

To analyse and locate long-term forest persistence, loss and gain, the historical forest map obtained above was intersected with the present-day forest map (BD Forêt® v2, https://cartes.gouv.fr/rechercher-une-donnee/dataset/IGNF_BD-FORET), produced by the IGN. BD Forêt® v2 is based on the photo-interpretation of aerial imagery acquired between 2005 and 2019, with a mean acquisition date of 2010. The planimetric accuracy of the boundaries is within ten metres for the outer contours of forest patches. Following the FAO (2000) definition, a forest patch was defined in this dataset as an area exceeding 0.5 ha, with an average width of at least 20 m, consisting of trees capable of reaching 5 m in height at maturity, and with a canopy cover greater than 10%. The definition also specifies that the area must not be primarily under agricultural or urban land use, thus excluding orchards (*e*.*g*., olive groves, and fruit-producing walnut and chestnut groves), as was already the case in the historical map. We retained plantations, including poplar plantations, in the present-day forest map. While the contemporary definition is based mostly on stand morphology (area, width, height and canopy cover) and secondarily on land-use (excluding land used for agricultural purposes), the 19^th^-century classification relied predominantly on the primary land product. To harmonise the two maps prior to their intersection, forest polygons smaller than the present-day minimum mapping unit of 0.5 ha were removed from the Ordnance map. Subsequently, polygons separated by less than 75 m were aggregated to eliminate small gaps within forested areas (e.g. forest roads and small openings) that are not represented in the present-day forest map. The first operation reduced the mapped forest area by 0.3%, whereas the second increased it by 1.2% relative to the original Ordnance forest map.

The intersection yielded four types of land-use transitions: (a) ancient forests, present in both the 19^th^-century and the present-day maps; (b) recent forests, which have developed since the 19^th^-century map; (c) deforested areas, present only on the 19^th^-century map; and (d) stable non-forest areas, which were unforested in both maps (Fig. 7).

**Fig. 7.**
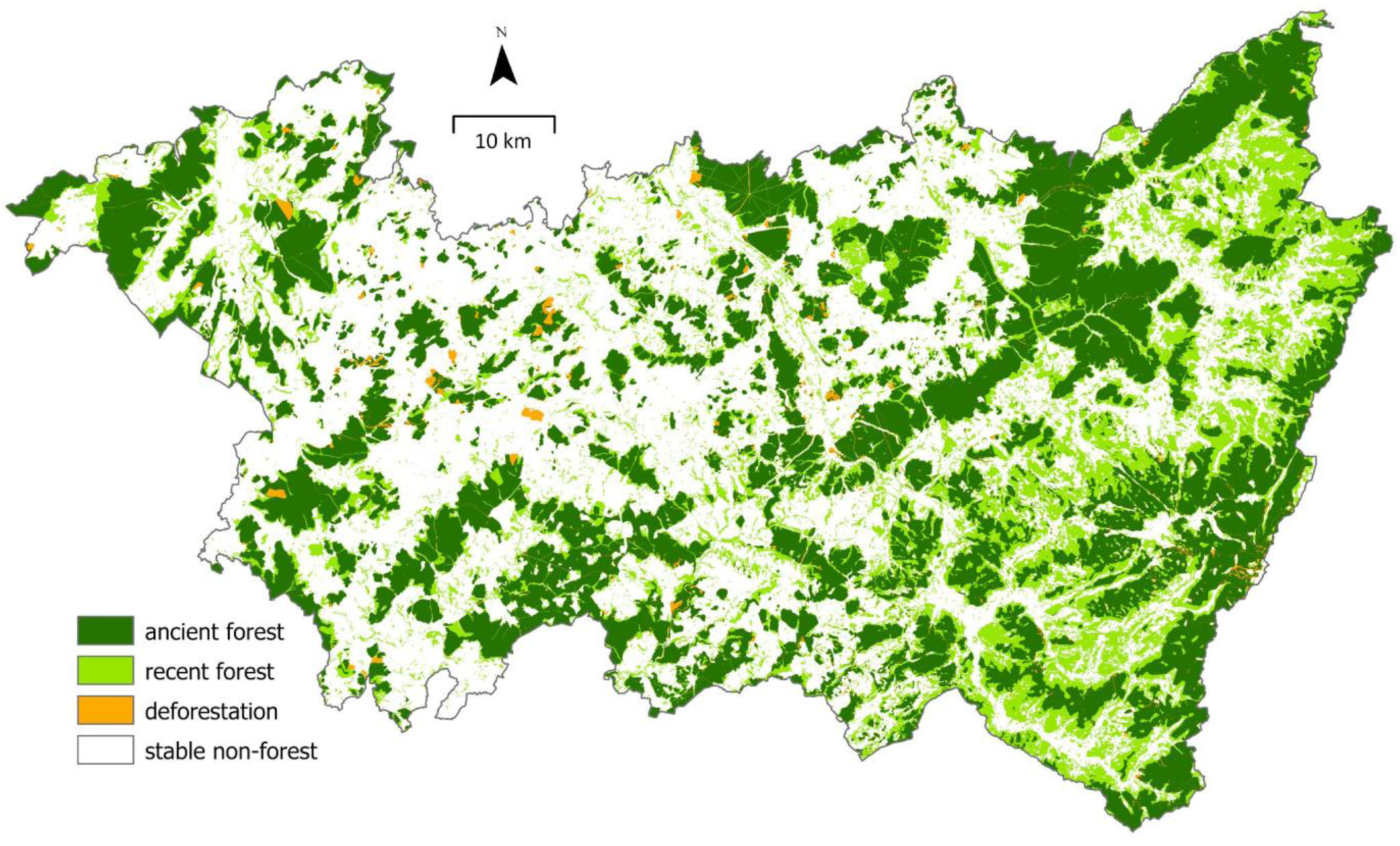
Forest cover dynamics in the Vosges department between the mid-19^th^ century and 2010. The map displays ancient forests (33% of department surface area), recent forests (19%), deforestation (1%), and stable non-forest areas (47%), obtained from the intersection of the Ordnance map (surveyed between 1826 and 1837 in this department) and the BD Forêt® v2 (2010).

Post-processing was performed to remove sliver polygons with an area of less than 1,000 m². These small features, generated by the intersection of the two layers, were primarily due to positional inaccuracies in the historical map and, to a lesser extent, in the present-day map. Thus, small polygons of ancient forest (0.1% of the total area of ancient forests), recent forest (0.1% of the total area of recent forests), deforestation (1.2% of the total deforested area) and stable non-forest area (0.03% of the total stable non-forest area) were reclassified according to the scheme presented in Appendix Table 3. While these reclassified polygons accounted for 36.7% of the total number of polygons, they represented only 0.08% of the total area. Therefore, this post-processing had virtually no effect on the areal metrics of the final map, while improving the accuracy of the boundaries and reducing the complexity and size of the vector layer.

## 3 Accuracy of the Ordnance map

Two factors can cause uncertainty in data quality: (a) the georeferencing process and (b) the reliability of information on historical land use.

### 3.1 Georeferencing process accuracy

We evaluated the influence of the georeferencing method and the number of control points on positioning error using a resampling procedure. For a sample of 28 map sheets in northern France (nine of which were also used in Section 2.3), we randomly selected subsets of control points of increasing size (10, 20, 25, 50, 100, 250, and 500) from the total pool available for each sheet. For each subset, we applied both a 6-parameter affine transformation and an elastic transformation (Delaunay triangulation followed by local 6-parameter affine transformations). Positioning RMSE was calculated for each method using the remaining out-of-sample points. This process was repeated 1,000 times for each subset size and each sheet. Finally, we calculated the arithmetic mean of these 1,000 RMSE values for each subset size and sheet combination (Fig. 8).

**Fig. 8.**
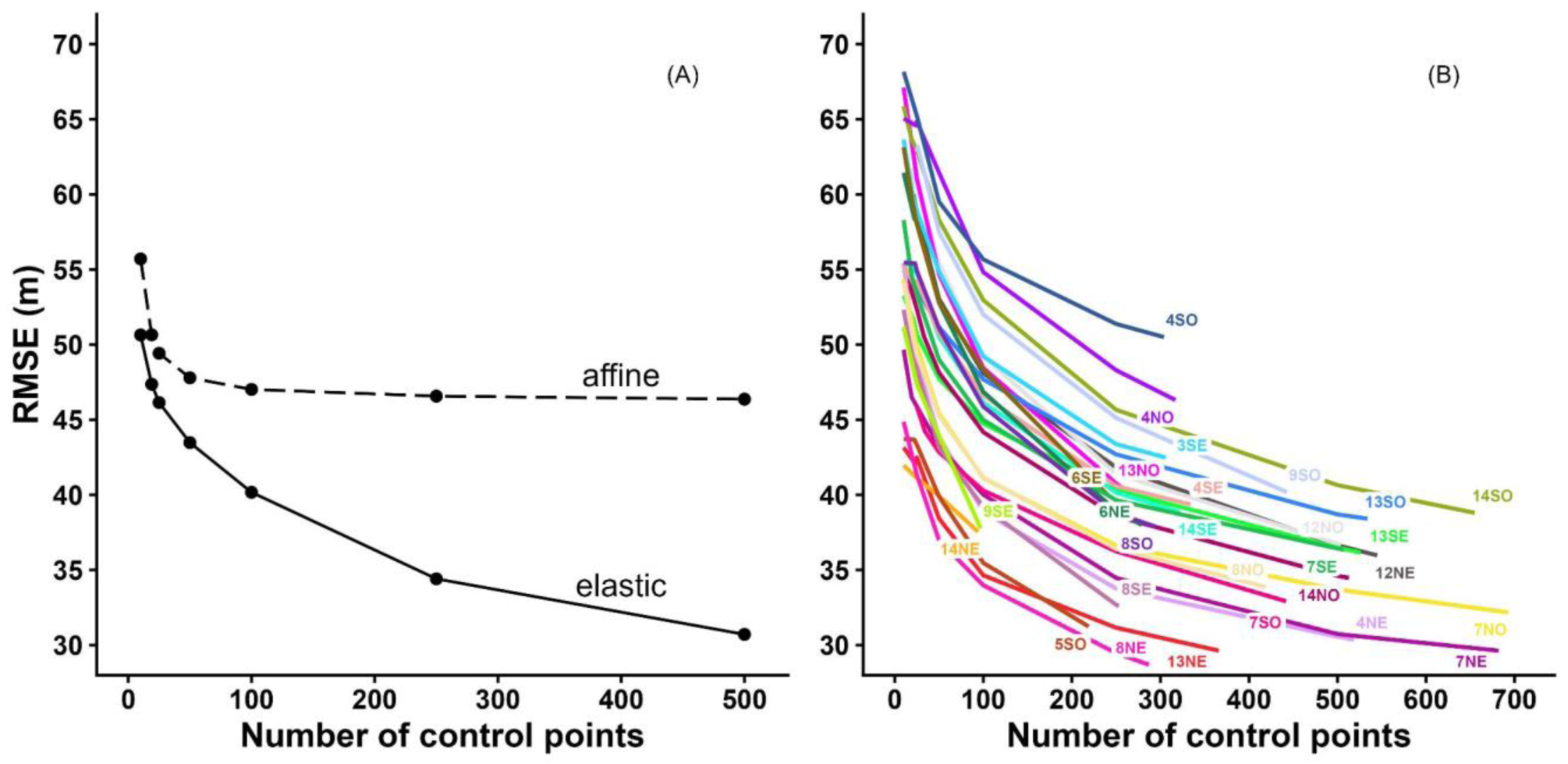
(A) Relationship between the positioning RMSE and the number of control points, comparing affine transformation (global method) and elastic transformation (local method) for the sheet 7NE (Arras). (B) Relationship between the positioning RMSE and the number of control points using the elastic transformation. Each line represents one of the 28 sheets tested.

The accuracy of georeferencing increased with the number of control points, regardless of the transformation method, up to a threshold of approximately 50 points (Fig. 8A). Beyond this threshold, the RMSE for the affine transformation reached an asymptote, showing no further significant reduction. In contrast, accuracy continued to improve when the number of control points increased with the elastic transformation. RMSE varied greatly between sheets (Fig. 8B). Despite its superior accuracy, elastic georeferencing was applied to a limited surface area (20.9% of the French territory, Table 2) due to the time required to identify high-quality control points and their scarcity in certain regions.

### 3.2 Reliability of land-use representation on the Ordnance map

To assess the quality of land-use representation on the Ordnance map, we compared it with the Napoleonic cadastre, a source considered to be more reliable (Rochel *et al*. 2017). Established during the same period, the cadastre is much more detailed: it records the use of every individual parcel and was established at scales ranging from 1:1,250 to 1:5,000, compared to the 1:40,000 scale of the Ordnance map. We compared the total area of each land-use category across 649 municipalities in the departments of Moselle (north-eastern France) and Vaucluse (south-eastern France) (Table 4).

**Table 4.** Comparison of historical land-use proportions between the Napoleonic cadastre and the Ordnance map, with associated statistics for 598 municipalities in the Moselle department (NE France) and 51 in the Vaucluse department (SE France). Data adapted from Rochel *et al*. (2017). Differences in percentage area between sources (Diff.) were assessed using paired-sample t-tests (*df*=597 for Moselle; *df*=50 for Vaucluse). Significance levels: NS not significant; * *p* < 0.05; ** *p* < 0.01; *** *p* < 0.001. *r* denotes Pearson’s correlation coefficient.

|  |  | Napoleonic<br>cadastre | Ordnance<br>map |  |  |
| --- | --- | --- | --- | --- | --- |
| Moselle (598 municipalities) |  |  |  |  |  |
| Survey date | Start | 1807 | 1828 |  |  |
|  | Average | 1825 | 1830 |  |  |
|  | End | 1852 | 1833 |  |  |
|  |  | Total area (%) |  | Diff. | *r* |
| Cultivated lands |  | 61.0 | 63.0 | +2.1 *** | 0.97 |
| Forests |  | 21.0 | 21.0 | +0.1 NS | 0.99 |
| Meadows |  | 9.5 | 10.5 | +1.0 *** | 0.93 |
| Built-up areas |  | 2.9 | 3.1 | +0.2 * | 0.80 |
| Vineyards |  | 1.3 | 1.5 | +0.1 ** | 0.99 |
| Pastures |  | 1.3 | 0.4 | -0.9 *** | 0.25 |
| Others |  | 3.0 | 0.4 |  |  |
| Vaucluse (51 municipalities) |  |  |  |  |  |
| Survey date | Start | 1809 | 1857 |  |  |
|  | Average | 1825 | 1859 |  |  |
|  | End | 1838 | 1861 |  |  |
|  |  | Total area (%) |  | Diff. | *r* |
| Cultivated lands |  | 40.6 | 48.1 | +7.6 *** | 0.94 |
| Forests |  | 25.7 | 26.1 | +0.3 NS | 0.95 |
| Pastures |  | 21.0 | 15.6 | -5.3 *** | 0.80 |
| Vineyards |  | 7.2 | 6.3 | -0.9 *** | 0.91 |
| Ponds and rivers |  | 1.7 | 1.0 | -0.7 *** | 0.83 |
| Meadows |  | 1.3 | 1.2 | 0.0 NS | 0.73 |
| Built-up areas |  | 0.3 | 0.7 | +0.3 *** | 0.73 |
| Others |  | 2.9 | 0.0 |  |  |

In both regions, the forest cover on the Ordnance map reflected that of the Napoleonic cadastre accurately. No systematic bias of over- or under-representation was observed (Rochel *et al*. 2017), and the correlation between forest cover percentages in the two sources was very high (r=0.99 in Moselle and r=0.95 in Vaucluse). The correlation between sources only fell below 0.8 for minor land-use categories: pastures in Moselle, and meadows and built-up areas in Vaucluse. Despite their limited extent in Moselle, vineyards were as well represented on the Ordnance map (r=0.99) as they were in Vaucluse (r=0.91). However, in both regions, croplands were over-represented on the Ordnance map compared to the cadastral data. While most land-use categories were represented by specific colours on the Ordnance map, croplands were indicated by the absence of colour. This may have led to a slight overrepresentation of cultivated areas, as any land-use types not explicitly recorded by the geographical engineers would, by default, have been merged into the “residual” cropland category.

Differences in scale, and the resulting degree of cartographic generalisation, might partly account for the variations between the two sources. In Vaucluse, the discrepancies in land-use coverage could also be attributed to a significantly longer time gap, 34 years on average, between the cadastral survey and the Ordnance map, compared to only 5 years in Moselle.

The strong agreement with the Napoleonic cadastre arose from the fact that forests often formed relatively compact blocks with stable boundaries, making them less sensitive to scale variations. Furthermore, woodland boundaries were systematically recorded on the cadastral maps, which were most often available as working documents for the geographical engineers during their field surveys for the Ordnance map. This confirms the value of the Ordnance map as a reliable source for delineating historical forests. However, in remote areas such as high-altitude mountains, forest representation is occasionally inaccurate; forests are sometimes depicted at elevations that are too high. Conversely, small woodland patches are sometimes omitted from historical maps (Everard et al. 2025), and from the Ordnance map in particular (Lathuillière & Gironde-Ducher 2021). This contributes to an underestimation of ancient forest extent and highlights the need to avoid relying solely on historical maps when identifying such forests, and to draw, wherever possible, on the full range of tools available in history, archaeology and palaeoecology (Everard et al. 2025, Bürgi & Östlund 2026).

## 4 A new estimate of the forest transition minimum

Forest area almost doubled (+81%) between 1843 and 2010, increasing from 18.1% to 32.7% of the total land area (Fig. 9), with large geographic variations (Fig. 10A-C and Appendix Table 5). In 1843, forest cover was generally lower in western France and higher in the east, with the exception of a few departments in the southwest that were more forested (departments 24 and 40). Present forest cover is mainly higher south of a line joining south-western to north-eastern France (departments 33 to 55 on Fig. 10B), reaching its highest levels in south-eastern France (departments 04, 06, 83), Corsica (2A, 2B) and the department of Landes (40). Consequently, forest cover increased more sharply in southern France, south of a line between departments 33 and 74 (Fig. 10C).

**Fig. 9.**
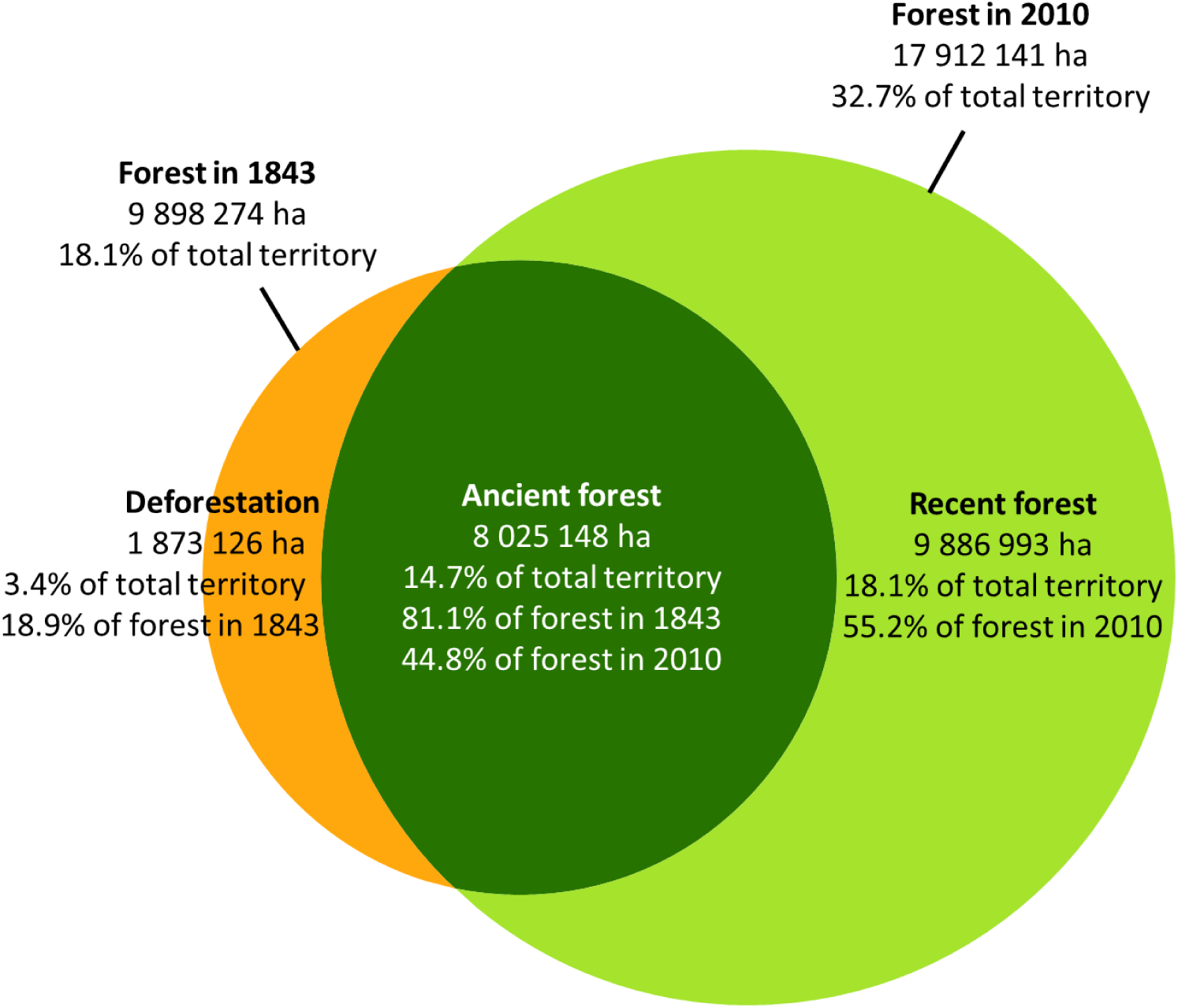
Surface area (ha) and shares (%) of historical forest (1843), present-day forest (2010), deforested areas, ancient forest and recent forest in France. Shares are provided relatively to the study area, the forest area in 1843 and/or the forest area in 2010, depending on the type of forest. The study area (‘total territory’, covering 547,215 km²) corresponds to the intersection between present-day French borders and the extent of the 19^th^-century Ordnance map. Shape areas are scaled proportionally to actual land surfaces.

**Fig. 10.**
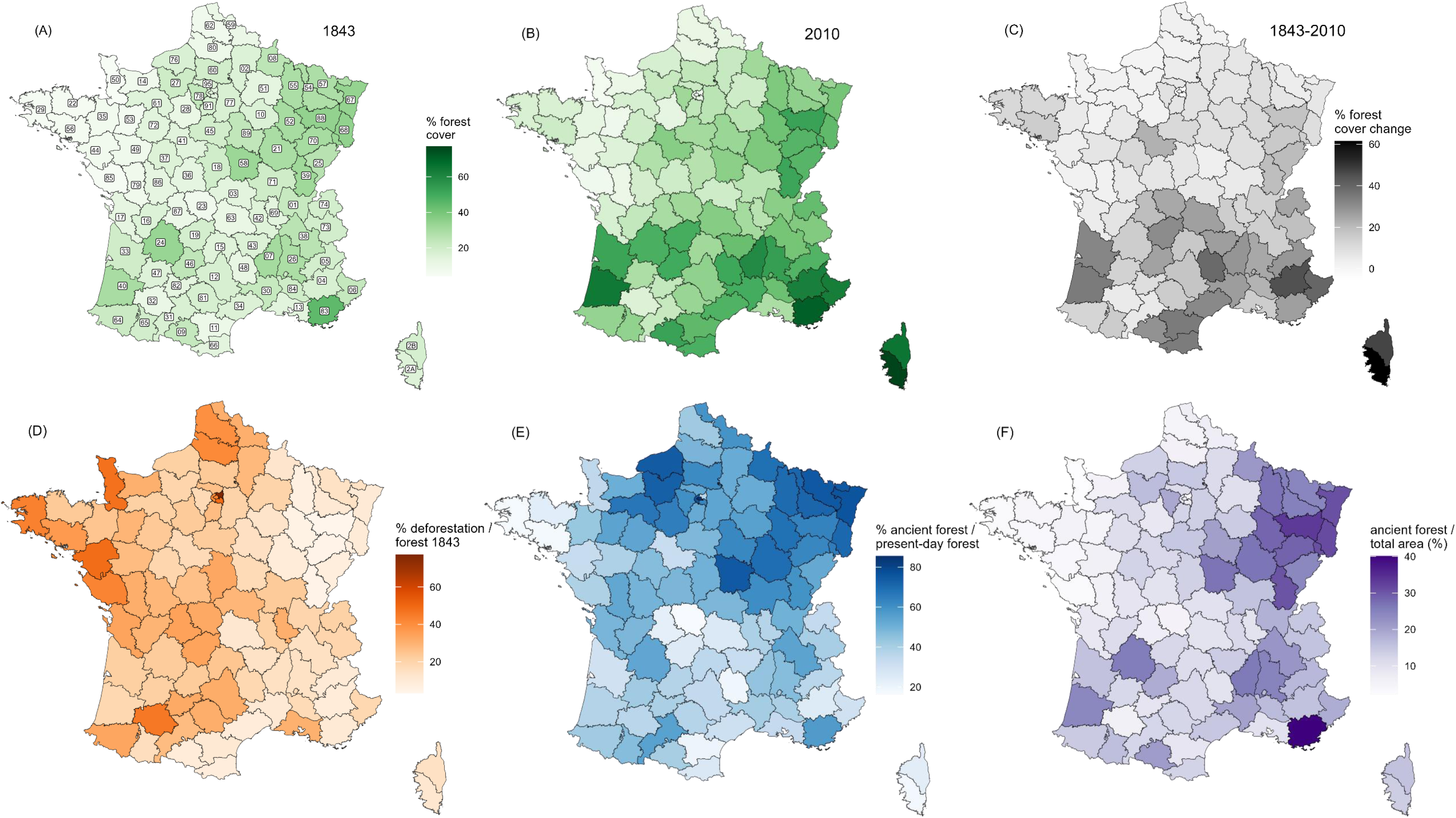
Forest cover dynamics by French department: (A) forest cover as a percentage of total area in 1843; (B) forest cover as a percentage of total area in 2010 (same colour scale as in A); (C) change in forest cover (percentage-point difference, 2010 minus 1843); (D) deforested area as a percentage of 1843 forest area; (E) ancient forest as a percentage of present-day forest area; and (F) ancient forest as a percentage of total area.

We hereby provide an upward revision of the forest area during the forest transition minimum to 9.9 million ha. Cinotti (1996) previously estimated this minimum at 9.2 ± 0.3 million ha for 1830, based on national aggregate statistics published by the Ministry of Agriculture and the Napoleonic cadastral services. We believe that a comparative department-level analysis of the areas derived from our new map versus those reported in the 19^th^-century cadastral statistics could yield further valuable insights.

Recent forests account for the majority (55.2%) of the present-day total forest area. Present-day forest cover is the net sum of three components: the initial forest area in 1843, the area of recent forests and the (negative) area of deforestation. We analysed the relative contribution of each of these three components to the variance in present-day forest cover across departments, using a simple variance decomposition. Notably, variations in present-day forest cover are driven more strongly by the expansion of recent forests since 1843 (accounting for 64% of the variance) than by the initial forest area existing in 1843 (37%). The spatial patterns and drivers of this expansion are analysed in Bontemps *et al*. (2020). Regarding the debate on the role of natural regeneration versus plantations, forest growth appears to be primarily driven by agricultural land abandonment and secondarily by intentional forest policies (Bontemps *et al*. 2020). In contrast, deforestation contributes negligibly to present-day inter-departmental variations in forest area.

Nonetheless, deforestation accounted for 18.9% of the forest cover in 1843, indicating that nearly one-fifth of these historical forests have disappeared over the last 170 years. This loss varied significantly among departments (Fig. 10D and Appendix Table 5), peaking in the most urbanised zones —particularly around Paris— and notably across the western half of France.

The proportion of ancient forest within the present-day forest cover averages 44.8% and exhibits strong spatial disparities (Fig. 10E and Appendix Table 5). The highest proportions are concentrated in north-eastern France (notably in the Ardennes, Burgundy, and Lorraine regions, in departments 08, 21, 52, 54, 55, 57, 58, 67 and 68) and Normandy (76, 27). In contrast, the lowest proportions are found in three zones: Brittany (22, 29, 56), the Massif Central (19, 23, 48, 87), and parts of the Mediterranean region (04, 11, 2A, 2B, 66). These three zones, which today contain a large proportion of recent forests, were often rich in heathlands, *garrigues*, or *maquis* at the time of the Ordnance map. Since then, these habitats have transitioned to forest. These differences are also partly attributable to shifts in forest definitions. Since the 19^th^-century classification was based on primary land use and associated products (*e*.*g*., grazing for heathland formations *versus* wood production for forests) rather than stand physiognomy, certain areas mapped, for instance, as *maquis* in Corsica during the 19^th^ century would likely be classified as forest under modern criteria (Panaïotis *et al*. 2017). More generally, the forested area in Mediterranean environments shown on the Ordnance map—and thus the extent of ancient forest identified by comparison with present-day maps—could be underestimated due to this conceptual discrepancy.

Whether forest expansion occurred linearly from the second half of the 19^th^ century to the beginning of the 21^st^ century, or through phases of acceleration and deceleration, remains to be further documented (but see Koerner *et al*. 2000; Bontemps *et al*. 2020). Importantly, instances of deforestation followed by reforestation between the Ordnance map and the present-day dataset are probably rare, as forest expansion was a widespread, continuous, and massive phenomenon across France throughout this period. We conducted a national-scale study to assess the proportion of so-called ancient forests—namely those present on both the 1843 and present-day maps—that were non-forested in 1950 (Appendix Table 6). We found that only 5.2% of ancient forests, according to the 1843-2010 comparison, were actually non-forested in 1950. In the Luberon region, Abadie (2018) analysed land use at an intermediate date (1958) using aerial photographs. Based on the analysis of a systematic 1 × 1 km grid (1,046 points in present-day forest), she showed that among forests classified as ’ancient’—*i*.*e*. appearing on both the Ordnance and present-day maps—6.7% were non-forested in 1958, having undergone a temporary conversion to non-forest land-use. Consequently, comparing the 1843 and 2010 states alone provides a reliable identification of ancient forests, *i.e.* those that have remained continuously forested. Nevertheless, mapping forests in the 1950s using the first national-scale aerial imagery represents the next step towards a better understanding of long-term forest dynamics (Gaubert *et al*., 2025). This would provide a snapshot of the forest at the onset of the agricultural ’Green Revolution’ following World War II. Another possible exception to forest continuity between 1843 and 2010 concerns areas temporarily destroyed during World War I and subsequently reforested. Fortunately, recent work has enabled the precise identification and mapping of these latter zones (Paradelle *et al*. 2023). In fact, the nature, intensity, and duration of the disturbances that interrupt forest continuity, as well as the time required for recovery, still require further investigation.

Ancient forest maps help identify priority areas for conservation and meet public policy requirements (Bergès and Dupouey 2021). For example, the Cévennes national park charter (2013) uses these data to identify zones where land clearing must be strictly limited and to target forest areas designated for non-intervention management. In its 2027-2042 management plan, the Ballons des Vosges Regional Natural Park identifies ancient and mature forests as core areas intended to form the backbone of a woodland habitat network. In addition, forest continuity is a criterion to be considered when identifying old-growth forests under the French National Action Plan for deadwood, senescent trees and subnatural forests. The map of ancient forest as a percentage of total territory (Fig. 10F) highlights French regions where ancient forest has become a scarce resource. At the European level, EU Directive 2023/2413 excludes biomass sourced from primary and old-growth forests from renewable energy targets and eligibility for financial support. Accurate identification of ancient and mature forests is therefore essential for compliance.

More generally, knowledge of this attribute of ancientness opens the door to a better understanding of present-day forests. Their species composition, biodiversity, soil quality, tree growth and water quality all depend on it, to name a few.

## 5 Access to the data & metadata description

Data are available at:

Map of 19^th^-century forests

DOI: https://doi.org/10.57745/KSCSQ2

Map of changes between 19^th^-century forests and present-day forests

DOI: https://doi.org/10.57745/JZ06QB

The complete dataset consists of 384 files, organised into two directories of 192 files each. The first directory contains the historical forest map, and the second contains the intersection between the historical and the present-day forest maps. In each directory, one map is provided per French administrative department (n=96). Each map is available in two alternative vector formats: a multi-file Shapefile (compressed in a .tar.gz archive) and an uncompressed GeoPackage (.gpkg file).

1. The first directory is named: forest_1843.

Each map contains three fields in its attribute table:

a. FOREST, coded as:

0: non-forest use;
1: forest use.
b. CODE_DEPT: French administrative department code.
c. area: surface area of the polygons (in m², using the Lambert 93 projection).

2. The second directory is named: forest_1843x2010.

Each map contains three fields in its attribute table:

a. LUT (land use trajectory) coded as:

AF: ancient forests (present in both the 19^th^-century map and the modern dataset).
RF: recent forests (established since the 19^th^-century map).
DEF: deforested areas (present in the 19^th^-century map only).
SNF: stable non-forest areas (unforested in both datasets).
b. CODE_DEPT: French administrative department code.
c. area: surface area of the polygons (in m², using the Lambert 93 projection).

File naming convention:

File names are composed of three parts separated by underscores:

1/ Map Type:

. FOR1843: forest layer derived from the Ordnance map (1843) or
. FOR1843x2010: forest-cover trajectory map generated by intersecting the historical Ordnance map (1843) with contemporary forest data (2010).
2/ Department code: D0XX (French administrative department).
3/ Version date: year and month of the version (e.g. 202512) to allow for future updates.

Example: FOR1843_D001_202512.tar.gz is the forest layer derived from the Ordnance map for department 01 (Ain), version produced in December 2025, in shapefile format, compressed as a .tar.gz file.

## 6 Funding

This work was partly financed by the FEDER Sylvalp program, a grant from the Ministry of the Environment, and the ‘Parc National des Pyrénées’.

## 7 Authors’ contributions

J.L.D., L.B., G.L., D.V., and A.G. conceived the project. Together with N.L., T.T., and T.L., they developed the methodology for map acquisition and georeferencing. The French National Institute of Geographic and Forest Information (IGN) provided the map scans. Data digitization was carried out primarily by IGN and INRAE, with contributions from all other authors. J.L.D., L.B., N.L., R.L. and M.B. conducted the initial analyses of the complete map of France. All authors reviewed and approved the final manuscript.

## 8 Acknowledgements

We would like to thank Baptiste Algoet, Matthieu Baconnet, Félix Benoît (⩾), Laurence Carnnot, Claire Crassous, Colette Favre, Vinciane Febvre, Marie Fougerouse, Sophie Giraud, Audrey Grel, Anna Hover, Sylvie Ladet, Jean-François Lopez, Arild Narbonne, Thomas Pauti, Mélanie Piroux, Jean-Baptiste Richard, Sylvain Rollet, Jean-Marie Savoie, Marie Thiberville, Rebecca Villemin, the students from AgroSup Dijon who worked within the framework of the Forests National Park, as well as Marie Bonnevialle, Maria Díaz de Quijano and Natacha Granger from the IPAMAC team for their assistance or involvement at various stages of this project.

This work was only possible thanks to the commitment and efforts of the IGN, and in particular the involvement of Edmond Alfamé, Francois Armandie, Isabelle Buisson, Christiane Cauchois, Régine Charlot, Françoise Crouzet, Dominique Donen, Jessy Fernandez, Julie Frohwirth, Bruno Fromont, Sylvie Geiger, Martine Gonthier, Marie-Catherine Goupil, Rémi Lefèvre, Valentine Lyot, Patrica Plard and Natacha Stavasius.

We sincerely thank GIP-Ecofor for their key role in this work, specifically for creating and managing the Cartofora website (http://www.gip-ecofor.org/cartofora/).

## 10 Appendix

Fig. 3. Areas of present-day France either outside the extent of the Ordnance map or lacking land-use data.

Fig. 6. Delineation of the areas of the Ordnance map that have been vectorised with level 4 georeferencing accuracy (mean error of 60 m) or level 5 (mean error of 34 m).

Table 3. Reclassification scheme for sliver polygons (below 1,000 m²) generated during the spatial intersection of historical and present-day forest maps.

Table 5. Area by land-use dynamics and department.

Table 6. Estimation of missed deforestation episodes between 1843 and 2010 in mainland France based on the analysis of 1950s aerial photographs.

## Notes

https://doi.org/10.57745/KSCSQ2

https://doi.org/10.57745/JZ06QB

### Competing Interest Statement

The authors have declared no competing interest.

### Summary of Updates

Addition of Appendix Figure 6 (map of areas vectorised at georeferencing accuracy level 4 or 5) and Appendix Table 6 (estimate of the percentage of so-called "ancient forests" that were actually deforested in the 1950s). Some other minor changes were made to the main text.

## References

Abadie J, Dupouey J-L, Avon C, Rochel X, Tatoni T, Bergès L (2018) Forest recovery since 1860 in a Mediterranean region: drivers and implications for land use and land cover spatial distribution. Landscape Ecology 33:289–305. doi.org/10.1007/s10980-017-0601-0

Abadie J, Dupouey J-L, Salvaudon A, Gachet S, Videau N, Avon C, Dumont J, Tatoni T, Bergès L (2021) Historical ecology of Mediterranean forests: Land-use legacies on current understorey plants differ with time since abandonment and former agricultural use. Journal of Vegetation Science 32:e12860. doi.org/10.1111/jvs.12860

Ågren AM, Lin Y (2024) A fully automated model for land use classification from historical maps using machine learning. Remote Sensing Applications: Society and Environment 36:101349. doi.org/10.1016/j.rsase.2024.101349

Andrieu D, Serres B (2024) L’intelligence artificielle apprend à lire le cadastre napoléonien [Artificial intelligence learns to read the Napoleonic cadastre]. Cartes & géomatique 255:73–83. doi.org/10.62437/cixz358cwvtgnjjlp-46087807

Auffret AG, Kimberley A, Plue J, Skånes H, Jakobsson S, Waldén E, Wennbom M, Wood H, Bullock JM, Cousins SAO, Gartz M, Hooftman DAP, Tränk L (2017) HistMapR: Rapid digitization of historical land-use maps in R. Methods in Ecology and Evolution 8:1453–1457. doi.org/10.1111/2041-210X.12788

Bacchus M, Dupuis J-C (1990) Une nouvelle carte de France par levé cadastral : bilan d’une idée révolutionnaire [A new map of France by cadastral survey: assessment of a revolutionary idea]. In Cartes, cartographes et géographes, Actes du 114^e^ Congrès National des Sociétés Savantes, Paris, 1989. CTHS 53-61

Bergès L, Avon C, Arnaudet L, Archaux A, Chauchard S, Dupouey J-L (2016) Past landscape explains forest periphery-to-core gradient of understorey plant communities in a reforestation context. Diversity and Distributions 22:3–16. doi.org/10.1111/ddi.12384

Bergès L, Feiss T, Avon C, Martin H, Rochel X, Dauffy-Richard E, Cordonnier T, Dupouey J-L (2017) Response of understorey plant communities and traits to past land use and coniferous plantation. Applied Vegetation Science 20:468–481. doi.org/10.1111/avsc.12296

Bergès L, Dupouey J-L (2021) Historical ecology and ancient forests: Progress, conservation issues and scientific prospects, with some examples from the French case. Journal of Vegetation Science 32(1):e12846. doi.org/10.1111/jvs.12846

Berthaut H (1898) La carte de France 1750-1898, Etude historique [The map of France 1750-1898, A historical study]. Service Géographique de l’Armée, 2 vol.

Bontemps J-D, Denardou A, Hervé J-C, Bir J, Dupouey J-L (2020) Unprecedented pluri-decennial increase in the growing stock of French forests is persistent and dominated by private broadleaved forests. Annals of Forest Science 77:98. doi.org/10.1007/s13595-020-01003-6

Bürgi M, Östlund L, Mladenoff DJ (2017) Legacy effects of human land use: Ecosystems as time-lagged systems. Ecosystems 20:94–103. doi.org/10.1007/s10021-016-0051-6

Bürgi M, Östlund L (2026) Bridging forest history and forest ecology: Why it is important to understand the history of forests and what are the main sources and methods. Ambio, 10 p. doi.org/10.1007/s13280-026-02502-y

Burst M, Chauchard S, Dupouey J-L, Amiaud B (2017) Interactive effects of land-use change and distance-to-edge on the distribution of species in plant communities at the forest-grassland interface. Journal of Vegetation Science 28:515–526. doi.org/10.1111/jvs.12501

Cateau E, Larrieu L, Vallauri D, Savoie J-M, Touroult J, Brustel H (2015) Ancienneté et maturité: deux qualités complémentaires d’un écosystème forestier [Ancientness and maturity: Two complementary qualities of forest ecosystems]. Comptes Rendus Biologies, 338(1):58–73. doi.org/10.1016/j.crvi.2014.10.004

Cinotti B (1996) Évolution des surfaces boisées en France : proposition de reconstitution depuis le début du XIX^e^ siècle [Evolution of forest cover in France: a proposed reconstruction since the early 19^th^ century]. Revue Forestière Française 48:547–562. doi.org/10.4267/2042/26776

Cousins SAO, Auffret AG, Lindgren J, Tränk L (2015) Regional-scale land-cover change during the 20^th^ century and its consequences for biodiversity. AMBIO 44:17–27. doi.org/10.1007/s13280-014-0585-9

Decocq G (coord.) (2022) Historical ecology: Learning from the past to understand the present and forecast the future of ecosystems. ISTE, 316 p. doi.org/10.1002/9781394169764

De Keersmaeker L, Onkelinx T, De Vos B, Rogiers N, Vandekerkhove K, Thomaes A, De Schrijver A, Hermy M, Verheyen K (2015) The analysis of spatio-temporal forest changes (1775-2000) in Flanders (northern Belgium) indicates habitat-specific levels of fragmentation and area loss. Landscape Ecol 30:247–259. doi.org/10.1007/s10980-014-0119-7

De Keersmaeker L, Roggemans P, Poelmans L, Priem F, Taillir S, Petermans T, Van Valckenborgh J (2026) Homogenization of Northern Belgian landscapes through centuries of reclamation, agricultural transition, and urbanization. Nature Communications. doi.org/10.1038/s41467-026-68594-y

Dupouey J-L, Bachacou J, Cosserat-Mangeot R, Aberdam S, Vallauri D, Chappart G, Corvisier de Villèle M-A (2007) Vers la réalisation d’une carte géoréférencée des forêts anciennes de France [Towards a georeferenced map of ancient forests in France]. Le Monde des Cartes 191:85–98.

Dupouey J-L, Bergès L (2026) A map of land-use trajectories generated by intersecting the historical Ordnance map with contemporary forest data (BD Forêt® v2, 2005-2019), Recherche Data Gouv, V1. <u>doi.org/</u>10.57745/JZ06QB

Dupouey J-L, Bergès L, Lafite R, Archaux F, Augé V, Bec R, Bellifa M, Bourguignon J, Buridant J, Burlin B, Caubet S, Chaléat A, Chauchard S, Cordonnier T, Decocq G, Delcamp M, Fleury J, Gaudin S, Gervaise A, Gautier G, Guilloux J-P, Hamel A, Heintz W, Janssen P, Labonne S, Lair P, Lallemant T, Landmann G, Larrieu L, Martin H, Michel C, Mollier S, Panaïotis C, Renaux B, Rochel X, Salvaudon A, Thomas M, Touzet T, Vallauri D (2026) The forest layer derived from the Ordnance map for mainland France, surveyed between 1818 and 1866, Recherche Data Gouv, V1. doi.org/10.57745/KSCSQ2

Eriksson S, Skånes H (2010) Addressing semantics and historical data heterogeneities in cross-temporal landscape analyses. Agriculture, Ecosystems & Environment 139:516–521. doi.org/10.1016/j.agee.2010.09.011

Everard A, Schofield JE, Mighall T, Timpany S, Plunkett G (2025) From stand-scale palaeoecology to the conservation of old-growth ‘ancient’ woodland. Ecological Indicators, 181, 114385. doi.org/10.1016/j.ecolind.2025.114385

FAO (2000) On definitions of forest and forest change. Forest Resources Assessment Programme, Working Paper 33, Food and Agriculture Organization of the United Nations, Rome, 14 p. http://www.fao.org/4/ad665e/ad665e00.pdf

Favre C, Audrey G, Granier E, Cosserat-Mangeot R, Bachacou J, Leroy N, Dupouey J-L (2017) Digitalisation des cartes anciennes. Manuel pour la vectorisation de l’usage des sols et le géoréférencement des minutes 1:40 000 de la carte d’Etat-Major [Digitizing historical maps: A manual for land-use vectorisation and georeferencing of the 1:40,000-scale Ordnance manuscript maps]. INRAE, v13.4, 61 p. Manuel_digitalisation_EM_Favre_et_al_2016_v12.8

Gaubert T, Bellet V, Hamrouni Y, Feuillet T, Birre D, Sheeren D (2025) Apprentissage par transfert entre images aériennes actuelles et historiques par un modèle génératif adapté à la segmentation sémantique du couvert arboré [Transfer learning between present-day and historical aerial imagery using a generative model adapted for semantic segmentation of tree cover]. Proceedings of the GRETSI’2025 Conference, Bordeaux, France, 1125–1128. gretsi.fr/data/colloque/pdf/2025_gaubert1742.pdf

Gobbi S, Ciolli M, La Porta N, Rocchini D, Tattoni C, Zatelli P (2019) New tools for the classification and filtering of historical maps. ISPRS International Journal of Geo-Information 8(10):455. doi.org/10.3390/ijgi8100455

Grabska-Szwagrzyk E, Jakiel M, Keeton W, Kozak J, Kuemmerle T, Onoszko K, Ostafin K, Shahbandeh M, Szubert P, Szwagierczak A, Szwagrzyk J, Ziółkowska E, Kaim D (2024) Historical maps improve the identification of forests with potentially high conservation value. Conservation Letters 17, e13043, 24 p. doi.org/10.1111/conl.13043

Helm A, Hanski I, Pärtel M (2006) Slow response of plant species richness to habitat loss and fragmentation. Ecology Letters 9:72–77. doi.org/10.1111/j.1461-0248.2005.00841.x

Herrault P-A (2015) Extraction de fragments forestiers et caractérisation de leurs évolutions spatio-temporelles pour évaluer l’effet de l’histoire sur la biodiversité : une approche multi-sources [Extraction of forest fragments and characterisation of their spatio-temporal changes to evaluate the effect of history on biodiversity: a multi-source approach]. PhD thesis, Toulouse le Mirail University, France, 397 p.

Herrault P-A, Sheeren D, Fauvel M, Paegelow M (2015) Vectorisation automatique des forêts dans les minutes de la carte d’état-major du 19^e^ siècle [Automated forest vectorisation in 19^th^-century Ordnance manuscript maps]. Revue Internationale de Géomatique 25:35–51. doi.org/10.3166/RIG.25.35-51

Jackson ST, Hobbs RJ (2009) Ecological restoration in the light of ecological history. Science 325:567–569. doi.org/10.1126/science.1172977

Janssen P, Bergès L, Fuhr M, Paillet Y (2019) Do not drop OLD for NEW: conservation needs both forest continuity and stand maturity. Frontiers in Ecology & the Environment, 17(7):370–371. doi.org/10.1002/fee.2086

Koerner W, Cinotti B, Jussy J-H, Benoît M (2000) Evolution des surfaces boisées en France depuis le début du XIXe siècle : identification et localisation des boisements des territoires agricoles abandonnés [Forest cover changes in France since the early 19th century: identification and localisation of afforestation on abandoned agricultural land]. Revue Forestière Française 52:249–269. doi.org/10.4267/2042/5359

Krauss J, Bommarco R, Guardiola M, Heikkinen RK, Helm A, Kuussaari M, Lindborg R, Öckinger E, Pärtel M, Pino J, Pöyry J, Raatikainen KM, Sang A, Stefanescu C, Teder T, Zobel M, Steffan-Dewenter I (2010) Habitat fragmentation causes immediate and time-delayed biodiversity loss at different trophic levels. Ecology Letters 13:597–605. doi.org/10.1111/j.1461-0248.2010.01457.x

Lalechère E, Jabot F, Archaux F, Deffuant G (2018) Projected regional forest plant community dynamics evidence centuries-long effects of habitat turnover. Journal of Vegetation Science 29 (3):480–490. doi.org/10.1111/jvs.12631

Lallemant T, Touzet T, Gervaise A (2017) Une méthodologie nationale pour le géoréférencement et la vectorisation des cartes d’état-major, minutes au 1/40 000 [A national methodology for the georeferencing and vectorisation of 1:40,000-scale Ordnance manuscript maps]. Revue Forestière Française 69(4-5):341–352. doi.org/10.4267/2042/67865

Lathuillière L, Gironde-Ducher M (2021) Méthode d’identification et de caractérisation des forêts anciennes grâce aux archives forestières [Method for identifying and characterising ancient forests using forestry archives]. Rendez-vous Techniques de l’ONF, 71-72:27-33. hal-03966002

Lindborg R, Eriksson O (2004) Historical landscape connectivity affects present plant species diversity. Ecology 85:1840–1845. doi.org/10.1890/04-0367

Martinez T, Hammoumi A, Ducret G, Moreaud M, Deschamps R, Piegay H, Berger J-F (2023) Deep learning ancient map segmentation to assess historical landscape changes. Journal of Maps 19(1), 2225071. doi.org/10.1080/17445647.2023.2225071

Mather, AS (1992) The forest transition. Area 24:367–379. doi.org/10.1016/0010-4485(92)90063-G

Mäyrä J, Kivinen S, Keski-Saari S, Poikolainen L, Kumpula T (2023) Utilizing historical maps in identification of long-term land use and land cover changes. Ambio 52:1777–1792. <u>doi.org/</u>10.1007/s13280-023-01838-z

Mollier S, Dupouey J-L, Kunstler G, Montpied P, Bergès L (2022a) Stronger legacy effects of cropland than of meadows or pastures on soil conditions and plant communities in French mountain forests. Journal of Vegetation Science 33:e13156. doi.org/10.1111/jvs.13156

Mollier S, Kunstler G, Dupouey J-L, Bergès L (2022b) Historical landscape matters for threatened species in French mountain forests. Biological Conservation 269:109544. doi.org/10.1016/j.biocon.2022.109544

Mollier S, Kunstler G, Dupouey J-L, Mulero S, Bergès L (2024) Forest management and former land use have no effect on soil fungal diversity in uneven-aged mountain high forests. Annals of Forest Science 81:2. doi.org/10.1186/s13595-023-01218-3

Navarro LM, Armstrong CG, Changeux T, Frisch D, Gil-Romera G, Kaim D, McClenachan L, Munteanu C, Szabó P, Baranov V, Blanco-Garrido F, Camarero JJ, García MB, Grace M, Izdebski A, Morueta-Holme N, Pando F, Schouten R, Spitzig A, Svenning J-C, Tribot A-S, Viana DS, Clavero M (2025) Integrating historical sources for long-term ecological knowledge and biodiversity conservation. Nature Reviews Biodiversity. doi.org/10.1038/s44358-025-00084-3

Panaïotis C, Barthet T, Vallauri D, Hugot L, Gauberville C, Reymann J, O’Deye-Guizien K, Delbosc P (2017) Carte d’état-major de la Corse (1864-1866). Occupation du sol et première analyse des forêts anciennes [The Ordnance map of Corsica (1864-1866): Land cover and preliminary analysis of ancient forests]. Ecologia mediterranea 43(1):49–64. doi.org/10.3406/ecmed.2017.2005

Paradelle N, Laslier M, Decocq G (2023) Automatic extraction of former WWI battlefields from ancient maps. In Neale CM, Maltese A (eds.), Remote Sensing for Agriculture, Ecosystems, and Hydrology XXV, Proc. of SPIE vol. 12727. <u>doi.org/</u>10.1117/12.2684009

Rochel X, Abadie J, Avon C, Bergès L, Chauchard S, Defever S, Grel A, Jeanmonod J, Leroy N, Dupouey J-L (2017) Quelles sources cartographiques pour la définition des usages anciens du sol en France[Cartographic sources for defining past land use in France]. Revue Forestière Française 69(4-5):353–370. doi.org/10.4267/2042/67866

Santana-Cordero AM, Szabó P, Bürgi M, Armstrong CG (2024) The practice of historical ecology: What, when, where, how and what for. Ambio 53:664–677. doi.org/10.1007/s13280-024-01981-1

Song D-X, Huang C, Sexton J, Channan S, Feng M, Townshend JR (2014) Use of Landsat and Corona data for mapping forest cover change from the mid-1960s to 2000s: Case studies from the Eastern United States and Central Brazil. ISPRS Journal of Photogrammetry and Remote Sensing 103:81–92. doi.org/10.1016/j.isprsjprs.2014.09.005

Stein ED, Doughty CL, Lowe J, Cooper M, Borgnis Sloane E, Bram DL (2020) Establishing targets for regional coastal wetland restoration planning using historical ecology and future scenario analysis: the past, present, future approach. Estuaries and Coasts 43:207–222. doi.org/10.1007/s12237-019-00681-4

Surasinghe T, Baldwin RF (2014) Ghost of land-use past in the context of current land cover: evidence from salamander communities in streams of Blue Ridge and Piedmont ecoregions. Canadian Journal of Zoology 92:527–536. doi.org/10.1139/cjz-2013-0307

Swetnam TW, Allen CD, Betancourt JL (1999) Applied historical ecology: Using the past to manage for the future. Ecological Applications 9:1189–1206. doi.org/10.1890/1051-0761(1999)009[1189:AHEUTP]2.0.CO;2

Szabó P (2015) Historical ecology: past, present and future. Biological Reviews 90:997–1014. doi.org/10.1111/brv.12141

Thomas M, Bec R, Abadie J, Avon C, Bergès L, Grel A, Dupouey J-L (2017) Changements à long terme des paysages forestiers dans cinq parcs nationaux métropolitains et le futur Parc National des Forêts de Champagne et Bourgogne [Long term changes in forest landscapes in five National Parks in metropolitan France and the future Champagne and Bourgogne National Forest Park]. Revue Forestière Française 69:387–404. doi.org/10.4267/2042/67868

Touzet T, Lallemant T (2016) Méthodologie nationale pour le géoréférencement et la numérisation des cartes d’État-major, minutes au 1:40 000 [A national methodology for the georeferencing and digitization of 1:40,000-scale Ordnance manuscript maps]. IGN, 178 p. docs.gip-ecofor.org/public/Rapport_Methodologique_IGN_v1-1.pdf

Vallauri D, Grel A, Granier E, Dupouey J-L (2012) Les forêts de Cassini. Analyse quantitative et comparaison avec les forêts actuelles [The Cassini forests: Quantitative analysis and comparison with present-day forests]. Rapport WWF/INRA, Marseille, 64 p. + CD. hal-01267936

Wulder MA, Roy DP, Radeloff VC, Loveland TR, Anderson MC, Johnson DM, Healey S, Zhu Z, Scambos TA, Pahlevan N, Hansen M, Gorelick N, Crawford CJ, Masek JG, Hermosilla A, White JC, Belward AS, Schaaf C, Woodcock CE, Huntington JL, Lymburner L, Hostert P, Gao F, Lyapustin A, Pekel J-F, Strobl P, Cook BD (2022) Fifty years of Landsat science and impacts. Remote Sensing of Environment 280:113195. doi.org/10.1016/j.rse.2022.113195

