## Appendix for "Maps of historical forests in France and their temporal continuity since the first half of the 19^th^ century"

**Fig. 3**. Areas of present-day France either outside the extent of the Ordnance map or lacking land-use data. These areas are situated in the Alps, along the south-eastern border (see Fig. 1 in the main text). Basemap source: Esri (2024), “World Topographic Map”, Available via ArcGIS Online.

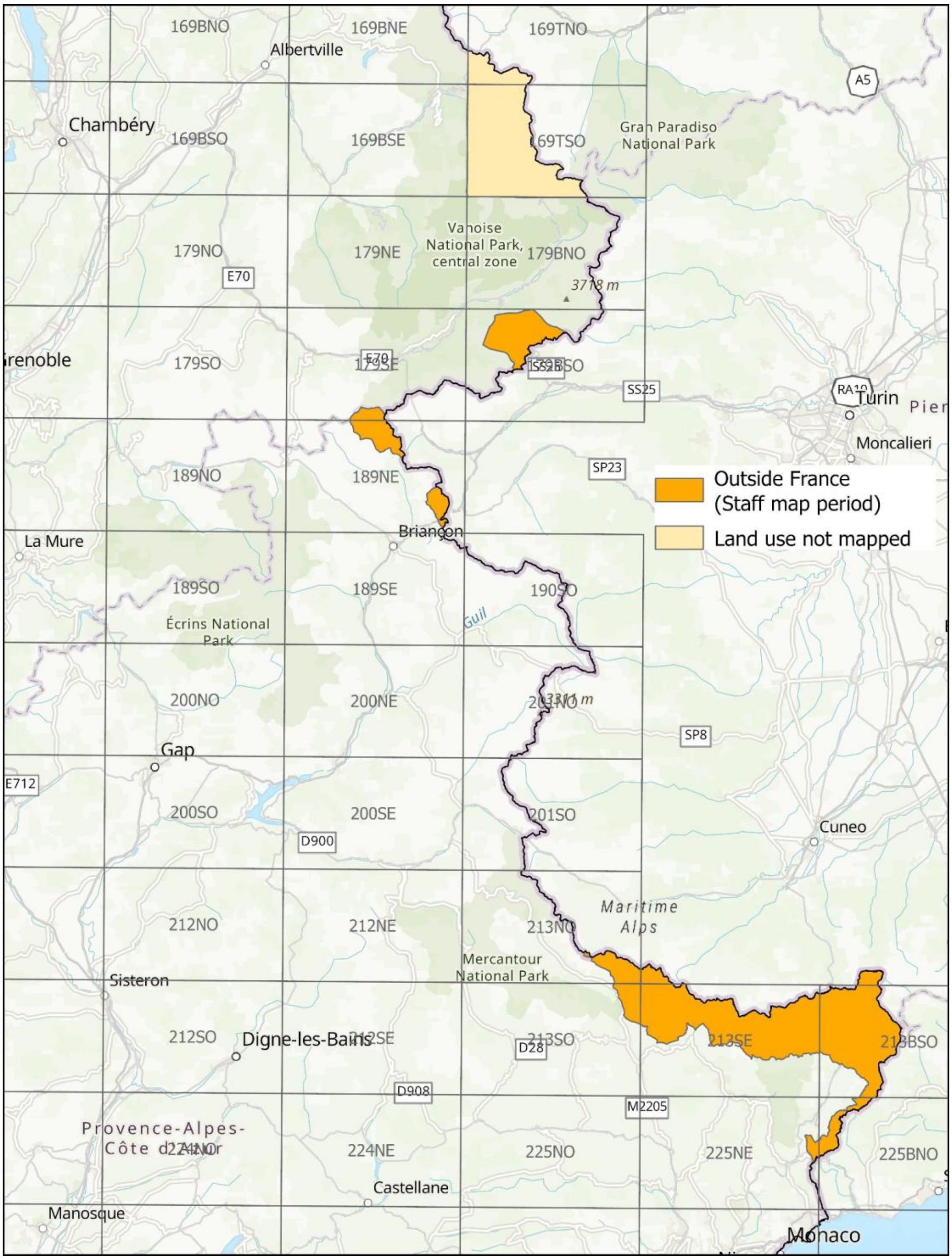

**Fig. 6**. Delineation of the areas of the Ordnance map that have been vectorised with level 4 georeferencing accuracy (mean error of 60 m) or level 5 (mean error of 34 m). These boundaries mostly follow the rectangular outlines of the original Ordnance map sheet grid (32 × 20 km) and, more rarely, those of current French administrative divisions (Pyrénées-Atlantiques and the Bauges Regional Natural Park in the Alps).

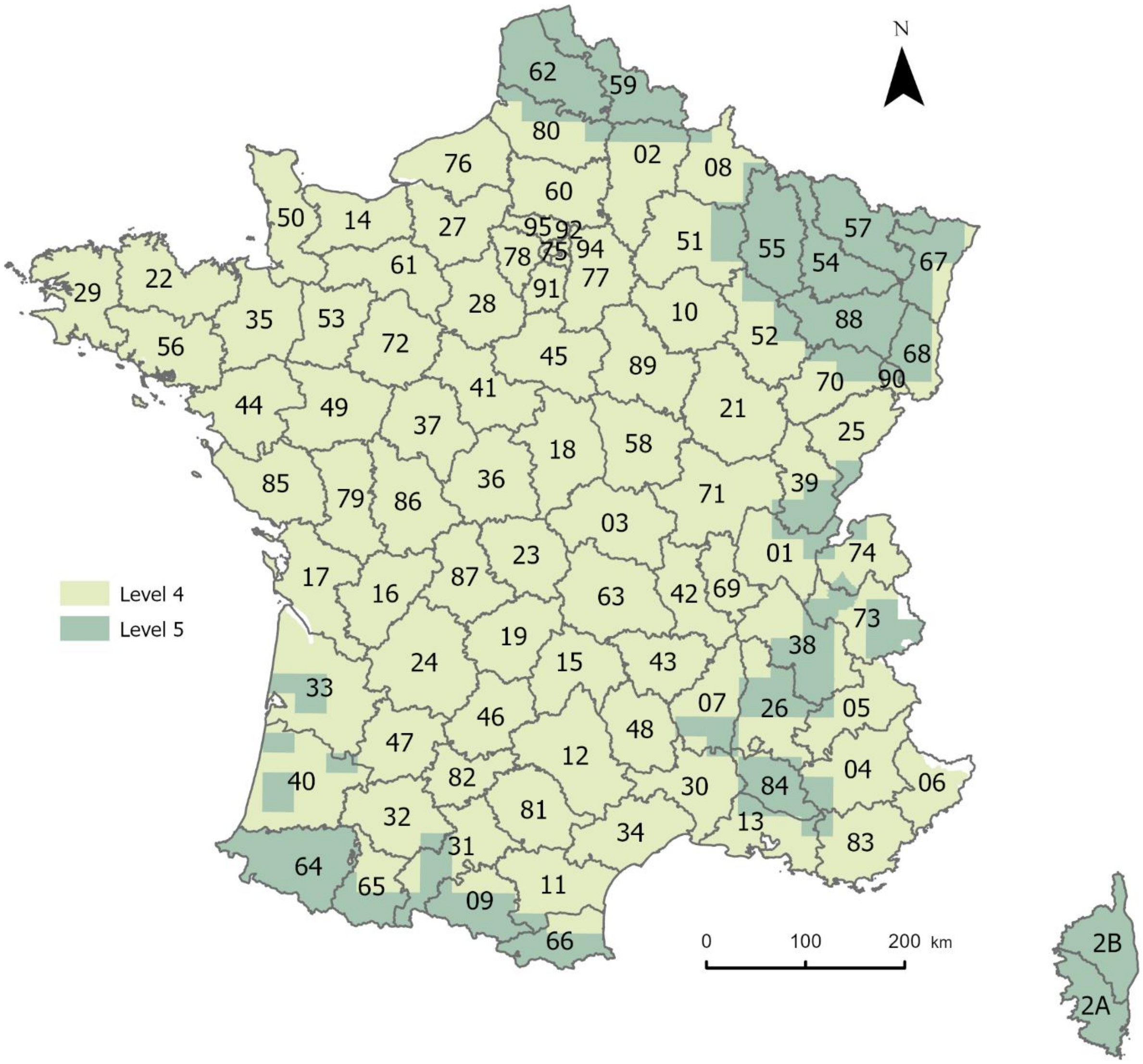

**Table 3**. Reclassification scheme for sliver polygons (below 1,000 m²) generated during the spatial intersection of historical and present-day forest maps. Each sliver was reclassified according to the class of its adjacent polygon(s). The rules applied here prioritise the present-day map, as its spatial accuracy is superior to that of the historical map. Combinations marked with a dash (–) represent non-existent cases. Following the application of these rules, certain polygon configurations may still result in some slivers requiring iterative reclassification; such rare cases were not further processed, thus leaving a small number of slivers in the final dataset.

| | **Sliver polygon category** | | | |
|---|---|---|---|---|
| **Adjacent polygons categories** | **stable non-forest area** | **recent forest** | **deforestation** | **ancient forest** |
| **stable non-forest area** | - | - | deforestation | - |
| **recent forest** | stable non-forest area | - | deforestation | ancient forest |
| **deforestation** | stable non-forest area | - | - | - |
| **ancient forest** | stable non-forest area | recent forest | deforestation | - |
| **stable non-forest area + recent forest** | - | - | stable non-forest area | ancient forest |
| **stable non-forest area + deforestation** | - | - | - | - |
| **stable non-forest area + ancient forest** | - | ancient forest | stable non-forest area | - |
| **recent forest + deforestation** | deforestation | - | - | recent forest |
| **recent forest + ancient forest** | stable non-forest area | - | deforestation | - |
| **deforestation + ancient forest** | deforestation | recent forest | - | - |
| **stable non-forest area + recent forest + deforestation** | - | - | - | recent forest |
| **stable non-forest area + recent forest + ancient forest** | - | - | stable non-forest area | - |
| **stable non-forest area + deforestation + ancient forest** | - | ancient forest | - | - |
| **recent forest + deforestation + ancient forest** | deforestation | - | - | - |

**Table 5**. Area by land-use dynamics and department. The total area for each department (the sum of the four columns) corresponds to the spatial intersection of the Ordnance map, the present-day forest map (BD Forêt® v2), and the department boundaries from the IGN BD TOPO® v2 map.

| Department | | Area (ha) | | | |
|---|---|---|---|---|---|
| Code | Name | Deforestation | Ancient forest | Recent forest | Stable non-forest |
| 01 | Ain | 25,350 | 104,981 | 104,612 | 343,538 |
| 02 | Aisne | 33,236 | 84,970 | 80,026 | 542,885 |
| 03 | Allier | 29,545 | 72,948 | 78,266 | 557,195 |
| 04 | Alpes-de-Haute-Provence | 10,555 | 109,468 | 325,671 | 253,468 |
| 05 | Hautes-Alpes | 10,365 | 95,549 | 167,163 | 289,423 |
| 06 | Alpes-Maritimes | 15,778 | 69,829 | 163,841 | 123,080 |
| 07 | Ardèche | 25,011 | 145,505 | 183,311 | 202,835 |
| 08 | Ardennes | 16,550 | 113,469 | 52,197 | 341,999 |
| 09 | Ariège | 10,025 | 102,849 | 153,263 | 224,970 |
| 10 | Aube | 13,085 | 84,907 | 68,703 | 436,082 |
| 11 | Aude | 6,784 | 56,216 | 225,686 | 345,029 |
| 12 | Aveyron | 48,217 | 105,829 | 201,624 | 521,674 |
| 13 | Bouches-du-Rhône | 21,658 | 49,945 | 97,225 | 335,468 |
| 14 | Calvados | 12,765 | 29,131 | 28,276 | 486,897 |
| 15 | Cantal | 21,018 | 64,775 | 116,765 | 374,873 |
| 16 | Charente | 27,955 | 67,040 | 71,810 | 430,571 |
| 17 | Charente-Maritime | 29,721 | 58,028 | 61,328 | 538,954 |
| 18 | Cher | 47,003 | 95,981 | 96,102 | 491,541 |
| 19 | Corrèze | 36,793 | 71,648 | 217,067 | 264,375 |

| Department | | Area (ha) | | | |
|---|---|---|---|---|---|
| Code | Name | Deforestation | Ancient forest | Recent forest | Stable non-forest |
| 2A | Corse-du-Sud | 7,301 | 55,223 | 254,387 | 84,614 |
| 2B | Haute-Corse | 12,495 | 73,093 | 234,898 | 150,089 |
| 21 | Côte-d'Or | 22,637 | 235,183 | 105,771 | 516,616 |
| 22 | Côtes d'Armor | 8,651 | 28,350 | 90,760 | 569,065 |
| 23 | Creuse | 13,952 | 29,170 | 149,764 | 367,056 |
| 24 | Dordogne | 69,931 | 237,281 | 208,075 | 406,972 |
| 25 | Doubs | 10,323 | 127,039 | 115,514 | 272,211 |
| 26 | Drôme | 27,028 | 162,880 | 186,985 | 279,020 |
| 27 | Eure | 23,954 | 98,377 | 36,890 | 440,308 |
| 28 | Eure-et-Loir | 14,941 | 49,918 | 27,092 | 501,237 |
| 29 | Finistère | 14,926 | 19,404 | 94,856 | 544,169 |
| 30 | Gard | 14,085 | 118,508 | 170,792 | 283,179 |
| 31 | Haute-Garonne | 29,494 | 78,823 | 64,733 | 462,758 |
| 32 | Gers | 31,065 | 36,697 | 64,168 | 498,100 |
| 33 | Gironde | 43,559 | 157,561 | 367,869 | 444,999 |
| 34 | Hérault | 8,633 | 91,636 | 203,197 | 318,873 |
| 35 | Ille-et-Vilaine | 11,988 | 36,562 | 47,953 | 585,350 |
| 36 | Indre | 23,440 | 66,679 | 70,978 | 528,854 |
| 37 | Indre-et-Loire | 18,605 | 78,923 | 91,383 | 426,864 |
| 38 | Isère | 29,810 | 172,520 | 144,988 | 440,672 |

| Department | | Area (ha) | | | |
|---|---|---|---|---|---|
| Code | Name | Deforestation | Ancient forest | Recent forest | Stable non-forest |
| 39 | Jura | 6,838 | 153,291 | 105,982 | 238,838 |
| 40 | Landes | 50,981 | 225,403 | 374,526 | 283,976 |
| 41 | Loir-et-Cher | 16,161 | 76,994 | 160,233 | 388,801 |
| 42 | Loire | 15,573 | 57,286 | 88,574 | 319,013 |
| 43 | Haute-Loire | 11,497 | 69,404 | 144,268 | 275,262 |
| 44 | Loire-Atlantique | 23,887 | 25,317 | 50,961 | 600,013 |
| 45 | Loiret | 29,707 | 100,698 | 108,977 | 441,877 |
| 46 | Lot | 28,881 | 97,656 | 163,030 | 233,043 |
| 47 | Lot-et-Garonne | 17,808 | 65,015 | 90,335 | 365,393 |
| 48 | Lozère | 11,379 | 50,371 | 212,180 | 243,601 |
| 49 | Maine-et-Loire | 15,937 | 42,634 | 67,855 | 590,842 |
| 50 | Manche | 11,879 | 13,054 | 24,076 | 547,957 |
| 51 | Marne | 21,464 | 85,727 | 80,500 | 631,938 |
| 52 | Haute-Marne | 9,956 | 177,989 | 82,488 | 355,180 |
| 53 | Mayenne | 9,864 | 23,986 | 20,116 | 467,390 |
| 54 | Meurthe-et-Moselle | 15,611 | 133,666 | 53,506 | 325,755 |
| 55 | Meuse | 11,612 | 167,991 | 72,697 | 371,140 |
| 56 | Morbihan | 14,240 | 24,700 | 116,716 | 526,669 |
| 57 | Moselle | 25,464 | 153,555 | 51,733 | 394,336 |
| 58 | Nièvre | 38,423 | 185,001 | 62,617 | 401,501 |

| Department | | Area (ha) | | | |
|---|---|---|---|---|---|
| Code | Name | Deforestation | Ancient forest | Recent forest | Stable non-forest |
| 59 | Nord | 17,893 | 40,499 | 28,320 | 488,976 |
| 60 | Oise | 26,354 | 86,444 | 54,264 | 422,049 |
| 61 | Orne | 19,708 | 73,007 | 36,291 | 485,597 |
| 62 | Pas-de-Calais | 20,158 | 31,079 | 42,650 | 575,468 |
| 63 | Puy-de-Dôme | 10,385 | 77,015 | 219,329 | 494,823 |
| 64 | Pyrénées-Atlantiques | 54,422 | 109,236 | 152,137 | 452,299 |
| 65 | Hautes-Pyrénées | 13,726 | 72,680 | 83,443 | 282,212 |
| 66 | Pyrénées-Orientales | 4,490 | 52,980 | 146,745 | 209,343 |
| 67 | Bas-Rhin | 16,141 | 147,445 | 46,501 | 269,644 |
| 68 | Haut-Rhin | 11,555 | 110,672 | 45,770 | 184,887 |
| 69 | Rhône | 13,932 | 31,613 | 53,466 | 226,867 |
| 70 | Haute-Saône | 6,684 | 155,442 | 94,095 | 282,837 |
| 71 | Saône-et-Loire | 39,583 | 132,742 | 85,674 | 603,407 |
| 72 | Sarthe | 20,733 | 67,135 | 58,324 | 478,378 |
| 73 | Savoie | 21,840 | 90,439 | 133,129 | 341,948 |
| 74 | Haute-Savoie | 17,919 | 68,103 | 131,423 | 241,768 |
| 75 | Paris | 696 | 904 | 153 | 8,792 |
| 76 | Seine-Maritime | 22,398 | 81,962 | 28,883 | 487,639 |
| 77 | Seine-et-Marne | 20,492 | 80,626 | 69,272 | 422,355 |
| 78 | Yvelines | 9,415 | 44,266 | 29,502 | 147,453 |

| Department | | Area (ha) | | | |
|---|---|---|---|---|---|
| Code | Name | Deforestation | Ancient forest | Recent forest | Stable non-forest |
| 79 | Deux-Sèvres | 14,701 | 32,870 | 29,102 | 527,370 |
| 80 | Somme | 23,923 | 34,212 | 36,213 | 524,071 |
| 81 | Tarn | 31,944 | 71,179 | 120,677 | 354,473 |
| 82 | Tarn-et-Garonne | 16,102 | 33,913 | 52,054 | 271,056 |
| 83 | Var | 26,927 | 241,479 | 184,299 | 149,106 |
| 84 | Vaucluse | 12,398 | 59,539 | 91,074 | 194,770 |
| 85 | Vendée | 14,642 | 19,802 | 31,073 | 608,089 |
| 86 | Vienne | 25,876 | 64,603 | 66,212 | 547,189 |
| 87 | Haute-Vienne | 19,677 | 37,575 | 128,400 | 370,108 |
| 88 | Vosges | 6,408 | 196,239 | 112,561 | 274,569 |
| 89 | Yonne | 26,422 | 152,530 | 93,316 | 473,923 |
| 90 | Territoire de Belfort | 1,342 | 18,790 | 10,110 | 30,828 |
| 91 | Essonne | 8,551 | 22,974 | 21,387 | 129,063 |
| 92 | Hauts-de-Seine | 1,160 | 1,722 | 547 | 14,132 |
| 93 | Seine-Saint-Denis | 2,380 | 697 | 1,264 | 19,349 |
| 94 | Val-de-Marne | 1,670 | 1,735 | 1,017 | 20,067 |
| 95 | Val-d'Oise | 5,085 | 14,367 | 12,957 | 92,932 |
| Total | | 1,873,126 | 8,025,148 | 9,886,993 | 34,936,225 |

**Table 6**. Estimation of missed deforestation episodes between 1843 and 2010 in mainland France based on the analysis of 1950s aerial photographs.

We sampled mainland France using the coordinates of a pre-existing soil monitoring network (Jolivet *et al.*, 2022). The plots of this network were established at the nodes of a systematic 16 × 16 km grid. A few points were shifted by up to 1 km during plot establishment.

We studied the 588 plots of this network whose land cover was classified as forest at the time of establishment (between 2000 and 2009). To characterize land cover in the 1950s, we used the earliest series of black-and-white aerial photographs available for the whole of France. These photographs were taken between 1945 and 1965. Their planimetric accuracy varies between 2 and 6 m. Within a 0.5-ha circular plot overlaid on the aerial photograph, we classified the land cover at each network point as forest or non-forest, according to the BD Forêt® v2 criteria.

| | | 1950s | | |
|---|---|---|---|---|
| | | forest | non forest | total |
| 1843 | forest | 308 | 17 | 325 |
| | non forest | 197 | 66 | 263 |
| | total | 505 | 83 | 588 |

Number of plots by land cover in 1843 and in the 1950s.
All studied plots were forested in 2000-2009.

From this table, we calculated the proportion of present-day forests that were classified as forest on the Ordnance map, in 1843, but went through a non-forested stage in the 1950s. These are considered "ancient forests" when only 1843 and 2010 are compared, but were actually temporarily deforested between these two dates. This proportion stands at 5.2% (17/(308+17)).

Reference

Jolivet C, Almeida-Falcon J-L, Berché P, Boulonne L, Fontaine M, Gouny L, Lehmann S, Maître B, Ratié C, Schellenberger E, Soler-Domingue N (2022) French Soil Quality Monitoring Network Manual. RMQS2: second metropolitan campaign, 2016–2027, Version 3, INRAE, Orléans, France. doi.org/10.17180/KC64-NY88
